# Transcriptome-wide meta-analysis of codon usage in *Escherichia coli*

**DOI:** 10.1101/2023.08.31.555696

**Authors:** Anima Sutradhar, Jonathan Pointon, Christopher Lennon, Giovanni Stracquadanio

## Abstract

The preference for synonymous codons, termed codon usage bias (CUB), is a fundamental feature of coding sequences, with distinct preferences being observed across species, genomes and genes. Accurately quantifying codon usage frequencies is useful for a range of applications, from guiding mRNA vaccine design, to elucidating protein folding and uncovering co-evolutionary relationships. However, current methods are either based on a single genome assembly, lack functional stratification, or are extremely outdated. To address this, we adopted a data-driven approach and developed Codon Usage Bias estimation from RNA-sequencing data (CUBSEQ), a fully automatic meta-analysis pipeline to estimate CUB at the trascriptome-level and for gene panels. Here, we used CUBSEQ to perform, to our knowledge, the largest and most comprehensive CUB analysis of the transcriptome and highly expressed genes in *Escherichia coli*, using RNA sequencing data from 6,763 samples across 72 strains. By capturing sequence variants of these genes through variant calls, we constructed a per-sample representation of the *E. coli* transcriptome revealing a rich mutational landscape. We then identified a set of 81 highly expressed genes with consistent expression patterns across strains, sample library size and experimental conditions, and found significant differences in CUB compared to transcriptome-wide genes and alternative codon usage tables. Finally, we found codons with a high relative frequency were often associated with a larger repertoire of isoaccepting tRNAs and not necessarily high tRNA abundance.

## Introduction

The degeneracy of the genetic code results in multiple codons being translated into the same amino acid. Nevertheless, these synonymous codons are not equally used, rather, they exhibit a nonuniform distribution, commonly known as codon usage bias (CUB), between organisms (interspecies codon usage), within the same genome (intraspecies codon usage) and even within genes [1, 2]. CUB is influenced by a complex orchestration of factors within and across genomes, including sequence features such as GC content [3], codon position [4], gene length [5], which impacts biological processes such as protein expression level [6], translation efficiency [7], and recombination rates such as biased gene conversion [8], as well as mRNA structure [9] and stability [10], and population size [11].

From an evolutionary point of view, CUB is thought to be largely driven by mutation rates, selection and genetic drift [12]. For example, codons in lowly expressed genes are thought to be largely influenced by mutation, whereas highly expressed genes (thought to be influenced by selection and mutation) have been found to have more biased synonymous codon usage in both Bacteria and Eukarya, likely to match the most abundant tRNAs and thus increasing translational efficiency and biological fitness [13]. This complements the finding that the use of a particular codon in a transcript causes later codons recognised by the same tRNA to be favoured (codon autocorrelation), thus increasing translation speed and fidelity through tRNA recycling [14].

The accurate quantification of codon usage benefits a wide range of applications, from fine-tuning gene expression for mRNA vaccine design [15], to understanding its relationship with protein folding and structure [16]. Furthermore, the process of back-translating amino acid sequences to their corresponding codons is a critical step for heterologous protein expression, having a significant impact on titers and process scale-up in bioreactors [17]. Choosing codons to mimic the CUB of the host organism can help to maximise protein yield and solubility. This process, often called codon optimisation (CO), is usually implemented by selecting the most frequent codon as a way to increase protein expression [18]; thus, obtaining accurate estimates of codon usage is pivotal to the success of most codon optimisation methods. As a result, understanding the rules underpinning codon usage will lead to greater insight into the processes controlling transcriptional and translational regulation, which in turn can be exploited for biotechnology applications.

Current and widely used codon usage databases, such as the Kazusa [19] and CoCoPUTs [20] codon tables, are calculated from a single reference genome of a given species, thus neglecting mutational processes pervading the transcriptome. The panel of genes is also another critical consideration in downstream calculations of CUB, but current codon usage tables are often non-selective and either use all genes or it is unclear how gene panels are selected. Moreover, CUB is usually quantified at the species level and not stratified by phenotype, for example the codon usage in highly expressed genes, thus limiting their utility for the biotechnology industry.

In this study, we tackle these challenges by employing a data-driven strategy that takes advantage of the vast amount of data derived from RNA sequencing studies [21]. We hypothesise that accurate codon usage estimates can be achieved by incorporating transcriptome variability. To accomplish this, we calculate codon relative frequencies based on a large collection of transcriptomes reconstructed from individual samples, enabling us to identify a robust panel of highly expressed genes. By harnessing the power of high-throughput RNA quantification assays, we aim to derive comprehensive insights into codon usage patterns associated with gene expression.

We developed the Codon Usage Bias estimation from RNA-sequencing data (CUBSEQ) pipeline (see Figure 1), which implements an end-to-end workflow to quantify gene expression, performs a transcriptome-based meta-analysis to build highly expressed gene panels, and then computes corresponding codon usage estimates. We then used CUBSEQ to process 6, 763 samples across 72 *E. coli* strains, a widely used model organism and workhorse protein expression system in the biotechnology industry, and compute robust estimates of codon usage at a transcriptome-wide level and in highly expressed genes.

**Figure 1:**
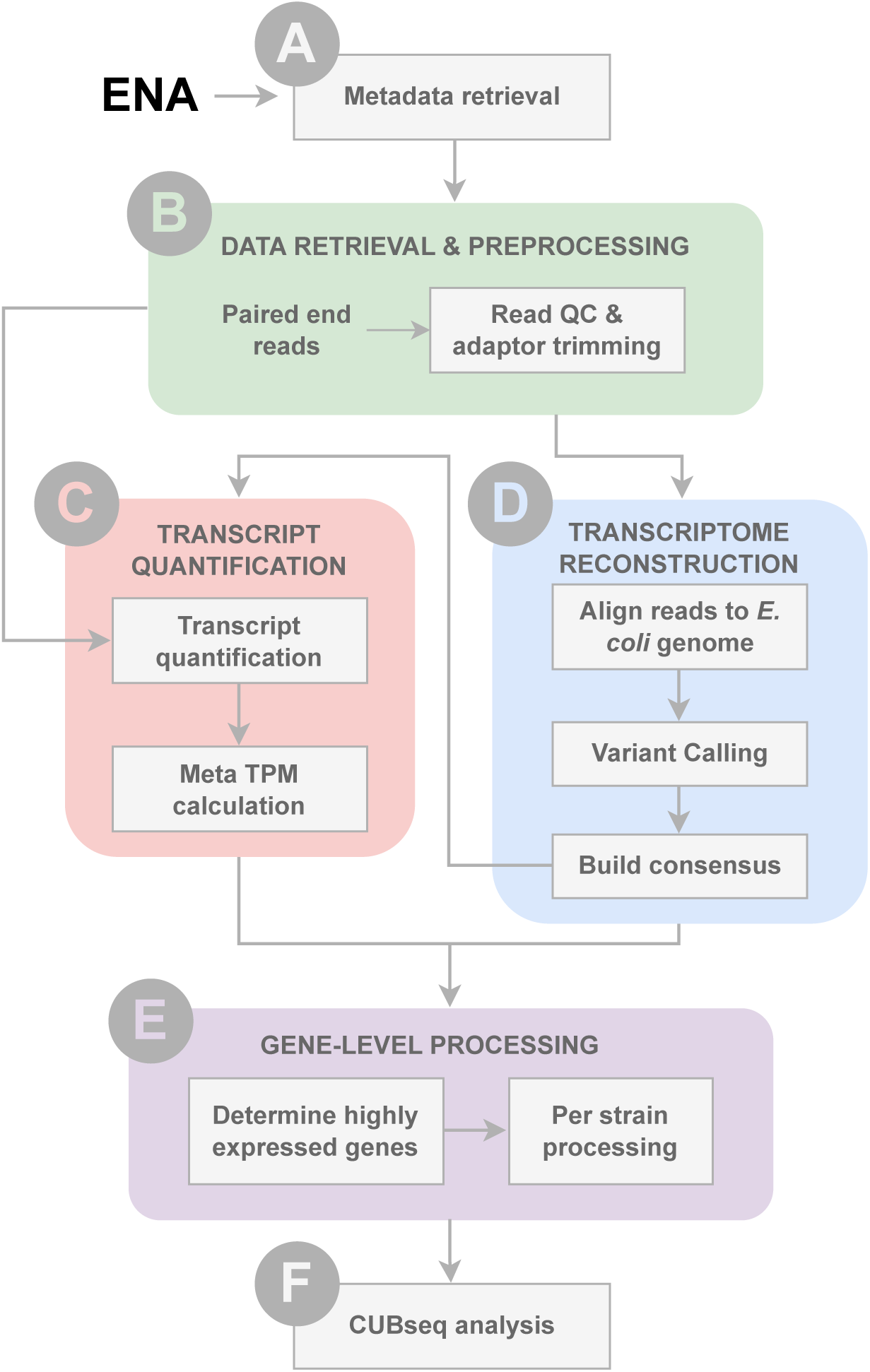
Schematic representation of the CUBSEQ pipeline. **(A)** The European Nucleotide Archive (ENA) is first queried to obtain metadata for *E. coli* transcriptomic reads from years 2012 to 2022, **(B)** out of which paired end fastq files are extracted and preprocessed to trim adaptors and filter out low quality reads (default parameters). Trimmed reads were then used to perform **(C)** alignment-free transcript quantification, which were then used to perform a meta-TPM calculation; and **(D)** transcriptome reconstruction by aligning to the *E. coli* K-12 reference genome, identifying synonymous variants and from this building a consensus sequence. **(E)** Meta TPM analysis results are then used to determine a panel of highly expressed genes, identified across all strains as well as on a per-strain level. **(F)** These results are then used for downstream codon usage analysis.

Our study found that while codon frequencies in *E. coli* are generally robust to mutational processes, they change dramatically when quantified with respect to highly expressed genes, suggesting an underlying functional stratification that can be exploited for biotechnology applications.

## Results

We developed a new transcriptome-based codon usage pipeline called CUBSEQ, and processed a total of 6, 763 runs generated by 507 studies from a total of 520 *E. coli* strains (or 72 taxonomic IDs) and produced over 17TB of data for samples sequenced between 2012 and 2022. Sequencing data came from 17 different types of Illumina machines and 13 different types of library selection methods. We processed a total of 95, 789, 815, 617 reads (14, 163, 805 average reads per sequencing run; see Supplementary Figure 1).

As this study involves processing thousands of samples, we performed principal component analysis (PCA) on the gene expression TPM dataset (see Methods) as a basic quality control step, to determine the sources of variation across sequencing runs. Similar to other meta-analysis studies, our analysis included incomplete or mislabelled metadata; nevertheless, we observed a clear clustering of taxonomy ID, library selection, instrument model, with limited variation observed in library size (see Supplementary Figure 4). As expected with *E. coli* samples, the majority of sequencing runs cluster together, with PC1 explaining 23.7% of the variation (see Supplementary Figure 5), however there are a clear group of sequencing runs (with large positive values across both PC1 and PC2) that cluster away from the majority and are likely driven by differences in strain, differing library selection of inverse rRNA selection and a smaller than average library size.

### *E. coli* transcriptome analysis reveals a rich mutational landscape

By integrating multiple experimental RNA-Seq samples we generated a comprehensive representation of the *E. coli* transcriptome at the sample level. To examine the mutational landscape of variants across the *E. coli* transcriptome, we performed variant calling on the alignments generated for each sequencing run using the *E. coli* K-12 MG1655 genome as reference.

Using a per-sample sequencing approach to construct a transcriptome-level genome, we reveal an extensive and diverse mutational landscape in the *E. coli* genome. Notably, when observing the allele frequency of alternate alleles for a particular variant across samples, we find that on average almost 40% of samples harbour alternate alleles across the transcriptome (Fig. 2A). A small proportion of individual loci were found to have very low alternate allele frequencies, which could be due to the variants being rare or residing in genomic regions difficult to capture by RNA-sequencing. The high read depth observed across alternate alleles (Supplementary Fig. 6B) increases confidence in the identification of true variants. Mean mapping quality (MQM) was also found to be high across all loci, indicating high confidence in alignment of reads in support of identified alternate alleles (Supplementary Fig. 6C).

**Figure 2:**
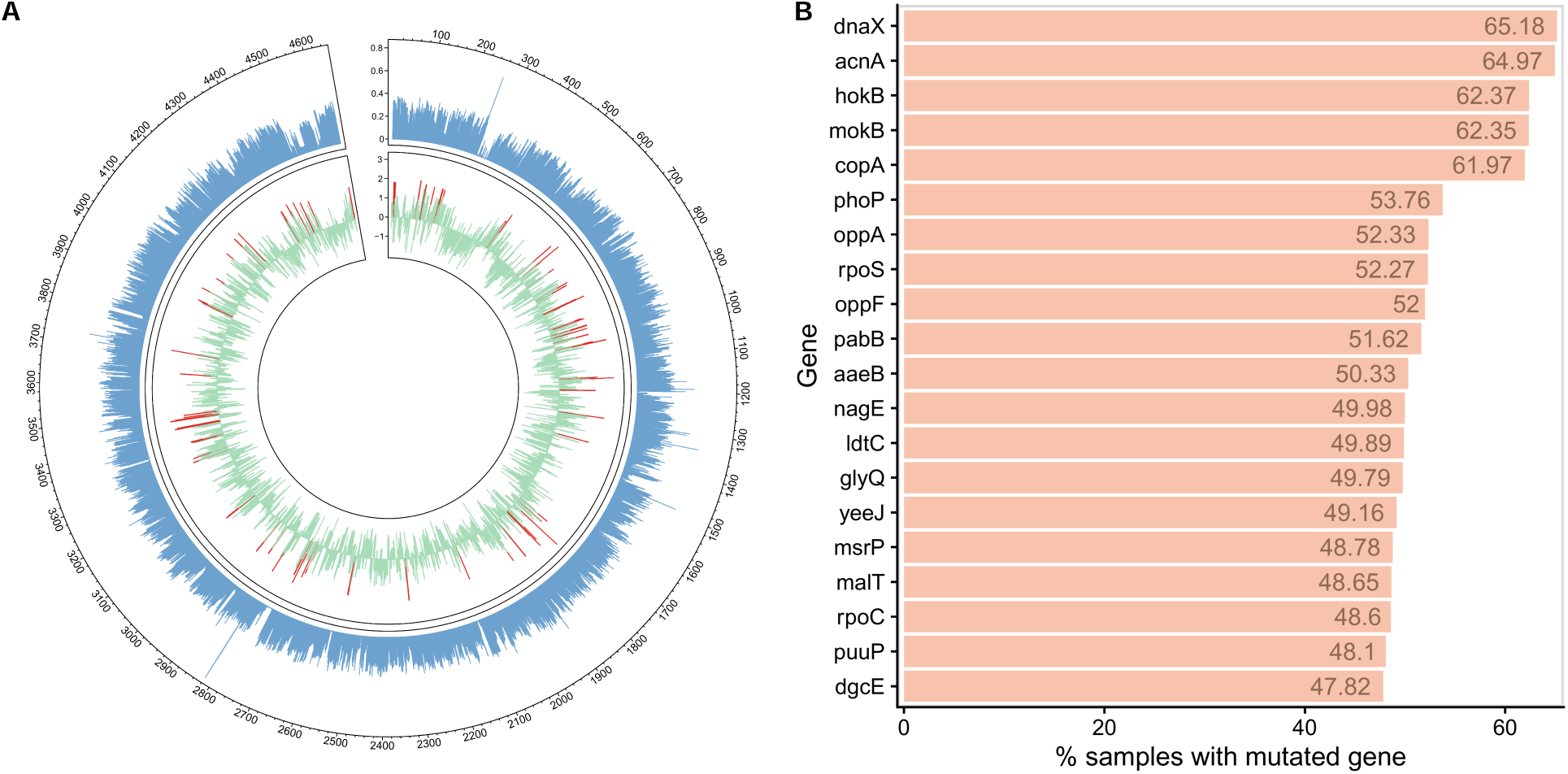
Mutational landscape of the *E. coli* transcriptome. **(A)** Circos visualisation of the transcriptome-level genome averaged across 6, 763 samples. The outer track shows the frequency of mutations per sample across loci (blue), determined by counting the total number of alternate alleles in called genotypes and normalising by the total number of samples. The inner track shows mean gene abundance scores for all transcriptome-wide protein-coding genes (green). Higher gene abundance scores indicate higher gene expression, with highly expressed genes indicated (red). **(B)** Genes (transcriptome-wide) found to harbour *≥* 1 alternate allele(s) in the highest proportion of samples, normalised across all 6, 763 sequencing runs.

In order to characterise the prevalence and distribution of genes carrying alternate alleles per sample, we then assessed the mutational landscape at the gene level, and determined the number of genes harbouring at least one alternate allele at any given loci across samples. We find the genes carrying at least one alternate allele across the majority of samples include *dnaX* and *acnA* (both mutated in 65% of sequencing runs), and *hokB*, *mokB* and *copA* (each mutated in 62% sequencing runs) (Fig. 2B). Both *dnaX* and *acnA* exhibit catalytic activity in DNA repair and during oxidative stress, respectively, making both genes susceptible to greater mutagenic pressures and possibly explaining the high frequency of mutations observed. *hokB* is also involved in the regulation of cell survival and stress response mechanisms, however could also be subject to additional adaptive selection pressures in response to its antisense antitoxin RNA (*SokB*). *CopA* is an interesting case, as its mRNA undergoes programmed ribosomal frameshifting to produce the first Heavy Metal-Associated (HMA) domain of the copA gene, serving as a cytoplasmic copper chaperone [22]. The high mutation frequencies observed therefore have the potential to affect the programmed ribosomal frameshifting mechanism, and could indicate the gene is under strong selective pressure.

Ribosomal protein *rpmJ* exhibited the presence of at least one alternate allele in the least number of samples (mutated in 0.23% of sequencing runs) (Supplementary Fig. 8B). Indeed, many genes encoding ribosomal proteins were found to carry at least one variant in a limited number of samples, suggesting that low coverage is not a confounding factor. Furthermore, this finding indicates the presence of stronger purifying selection pressures acting to select against potential synonymous mutations (as opposed to also non-synonymous mutations) relative to non-ribosomal genes.

Taken together, our study characterised a highly complex mutational landscape, which should be taken into account as part of any downstream codon usage analysis.

### CUBseq highly expressed genes are robust to strain variation

One of the key advantages of using bulk-processed RNA-sequencing data is the ability to obtain unbiased estimates of highly expressed genes across numerous samples and experimental conditions. In our study, we specifically focus on identifying high expression genes while excluding ribosomal genes, which are subject to unique selective pressures due to their involvement in translation machinery. This allows us to isolate and investigate other highly expressed genes that may hold significant potential for biotechnological applications, particularly in the context of codon optimisation for heterologous protein expression.

To achieve this, we use the CUBseq meta-analysis method to build a panel of highly expressed genes (HEGs) across all 6, 763 samples. Here we identified a panel of 81 genes, mostly involved in various metabolic processes, with 16 genes associated with macromolecule metabolic and organonitrogen compound metabolic processes (respectively), and 12 genes associated with cellular nitrogen compound biosynthetic processes (Fig. 3A).

**Figure 3:**
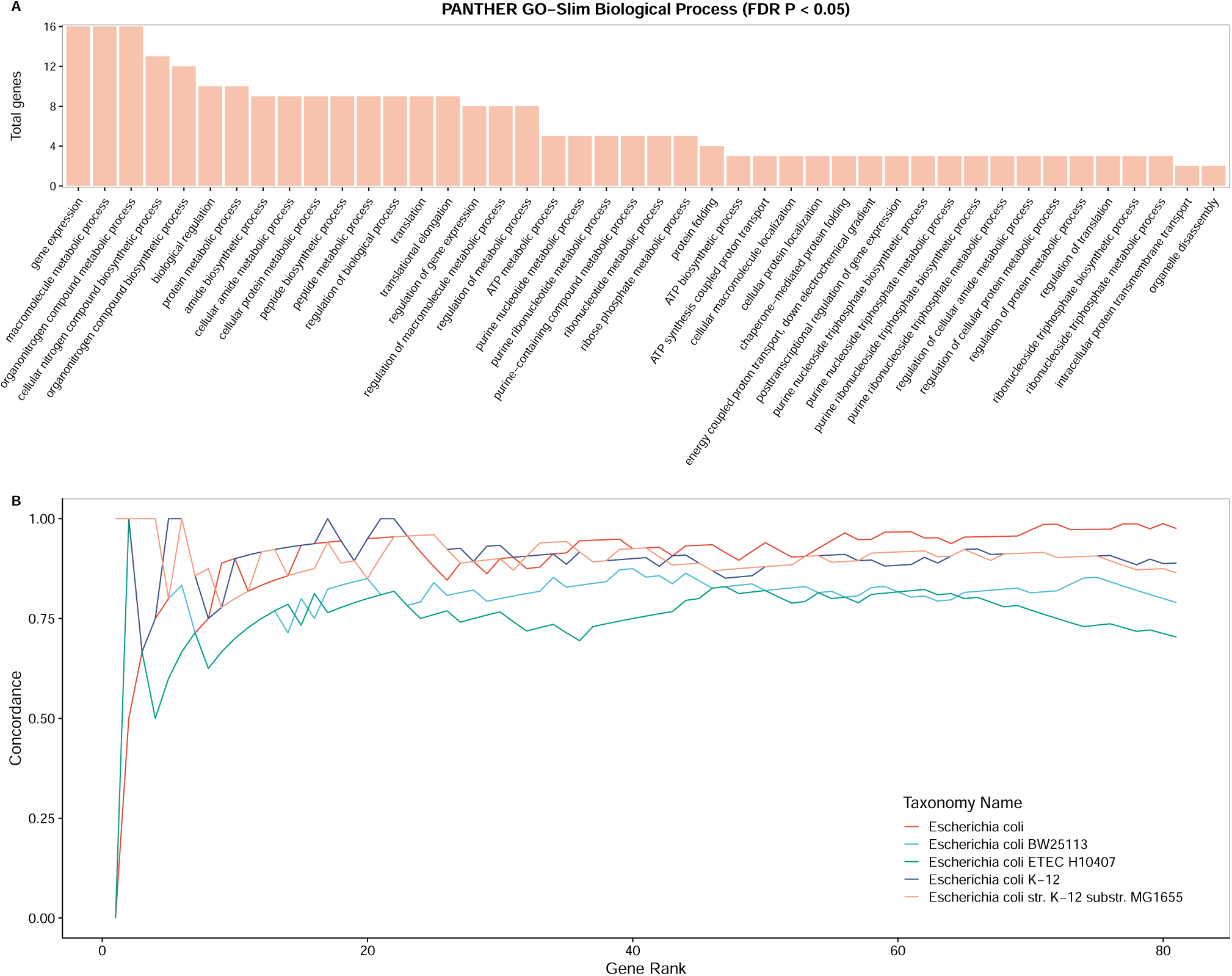
Transcriptome-wide panel of significantly highly expressed genes across the most common five *E. coli* strains show general stability in codon usage. **(A)** GO biological process terms from the PANTHER GO-Slim classification system, associated with our transcriptome-derived panel of 81 significantly highly expressed genes (filtered for FDR *P <* 0.05). **(B)** A concordance at the top (CAT) plot, showing concordance versus the rank of our 81 significantly highly expressed genes across the top five strains (i.e. strains having the greatest number of sequencing runs and representing 88% of the CUBseq dataset). The different colours represent different *E. coli* strains, identified by taxonomy ID. The highly expressed genes for each strain are compared to our transcriptome-based panel of highly expressed genes as reference, that is, all the strains combined. Note that *”Escherichia coli”* does not represent these combined strains, but instead the most frequent taxonomy ID (tax id 562) provided in the metadata.

To determine robustness of our HEG panel across strains, we performed a concordance at the top (CAT) analysis (Fig. 3B) comparing high expression gene rankings across the top five most represented strains (representing 87.73% of the entire dataset) to the genes derived from the meta-analysis (i.e. all *E. coli* strains combined) as reference. It is important to note that these sequencing runs originate from hundreds of different experiments, and so we would not expect strains to reach 100% concordance. However, when comparing gene rankings of the 5 most represented strains, we find our set of HEGs rapidly gain concordance regardless of strain and remain relatively stable as gene rank increases. As expected, we find the top taxa for *E. coli* to show the strongest concordance (reaching final concordance of 97.53%), followed by *E. coli* K-12 (90.12%), *E. coli* sub-strain MG1655 (86.42%), *E. coli* BW25113 (79.01%) and *E. coli* ETEC H10407 (70.37%). In total, all top five strains reach over 70% concordance, suggesting our HEG panel is a bona-fide set of highly expressed *E. coli* genes for robust analyses of strain-wide codon usage patterns.

### CUBseq highly expressed genes exhibit stronger codon usage bias compared to transcriptome-wide genes

To compare how codon usage bias differs in CUBseq transcriptome-wide genes versus highly expressed genes, we calculated codon relative frequencies for the respective gene panels (Fig. 4, 5). Methionine and Tryptophan were excluded from this analysis, as both are encoded by one codon.

**Figure 4:**
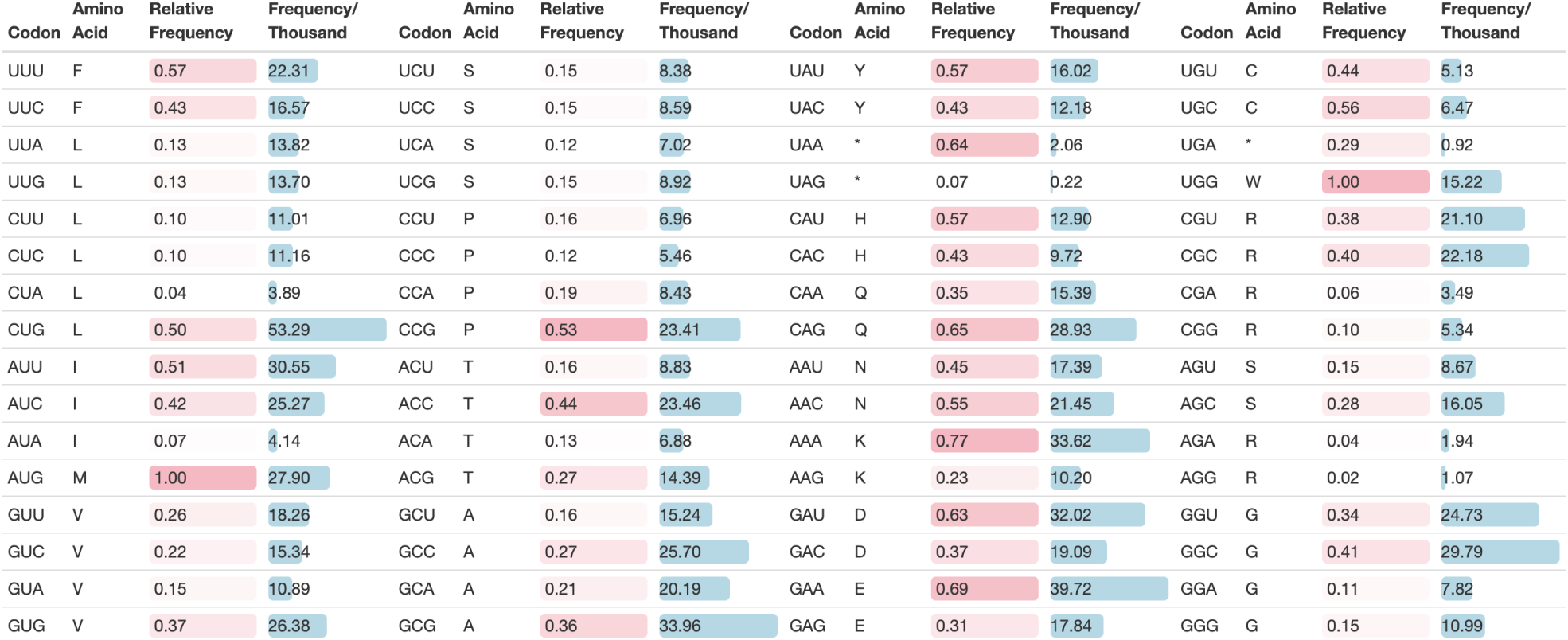
Codon usage frequencies based on transcriptome-derived frequencies of all protein-coding genes in *E. coli*. The codon usage frequencies are calculated based on 8, 968, 928, 288 total codons. Values are given as “Relative Frequency” (i.e. as a percentage relative to the respective amino acid), or as Frequency per Thousand (where total codon count for a particular codon is divided by the sum of all codons, then multiplied by a thousand). Relative frequencies are coloured by pink tiles, with higher frequency values being depicted by darker shading. Frequency per thousand values are depicted with blue bars, with higher frequencies having a greater bar length.

**Figure 5:**
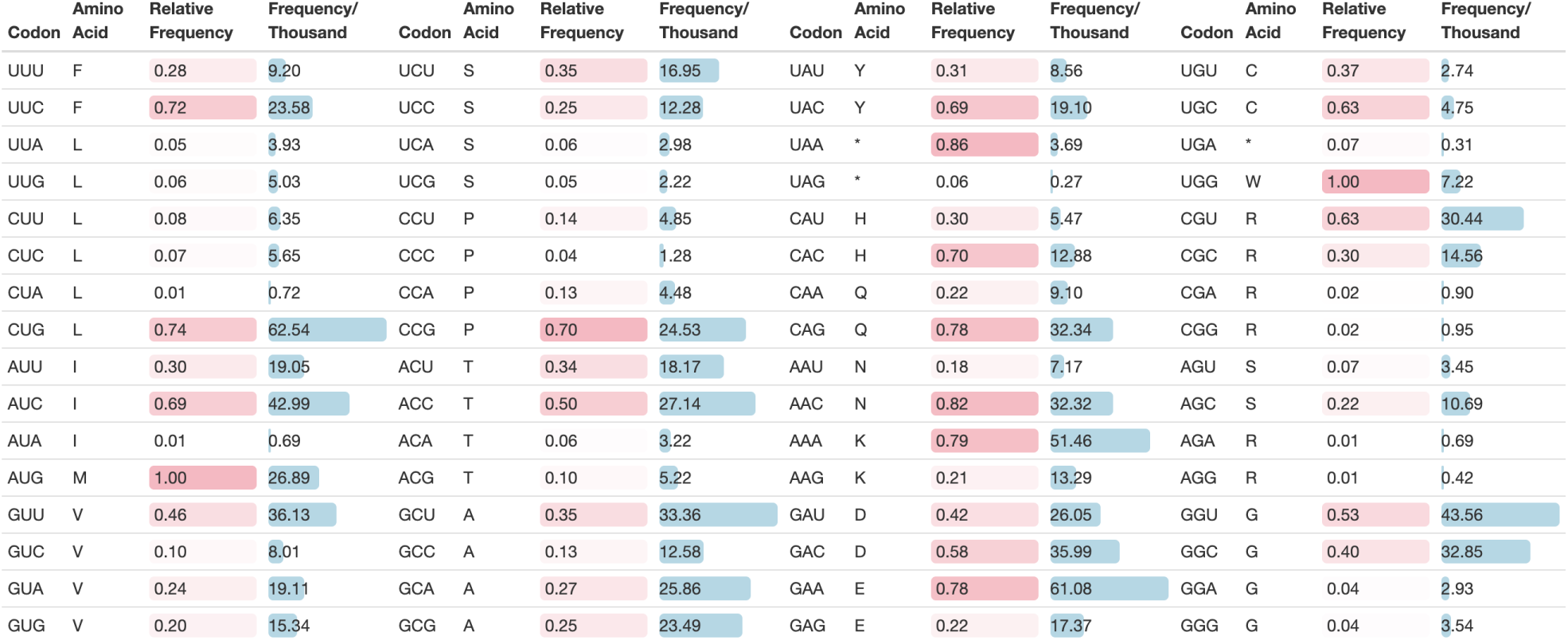
Codon usage frequencies, based on the transcriptome-derived panel of 81 significant highly expressed protein-coding genes in *E. coli*. The codon usage frequencies are calculated based on 128, 314, 399 total codons. Values are given as “Relative Frequency” (i.e. as a percentage relative to the respective amino acid), or as Frequency per Thousand (where total codon count for a particular codon is divided by the sum of all codons, then multiplied by a thousand). Relative frequencies are coloured by pink tiles, with higher frequency values being depicted by darker shading. Frequency per thousand values are depicted with blue bars, with higher frequencies having a greater bar length.

We first investigated whether distinct signatures of codon usage were present. Specifically, we assessed codon preference at the amino acid level by computing the odds of observing the most frequent codon compared to other synonymous codons. Within the set of transcriptome-wide genes, we observed Lysine to have the strongest preference for its most frequent codon, with a preference ratio of 3.3 : 1 in favour of the *AAA* codon over the *AAG* codon. This was followed by Glutamate, having a preference ratio of 2.2 : 1 in favour of the *GAA* codon over the *GAG* codon. Serine had the lowest preference ratio of 0.39 for its *AGC* codon over its five other synonymous codons; followed by Alanine and Valine having a preference of 0.56 and 0.59 for their most frequent codons *GCG* and *GUG*, respectively. Overall the preference for the codons of transcriptome-wide genes was found to be relatively weak, averaging at 1.23 across amino acids (Fig. 6A).

**Figure 6:**
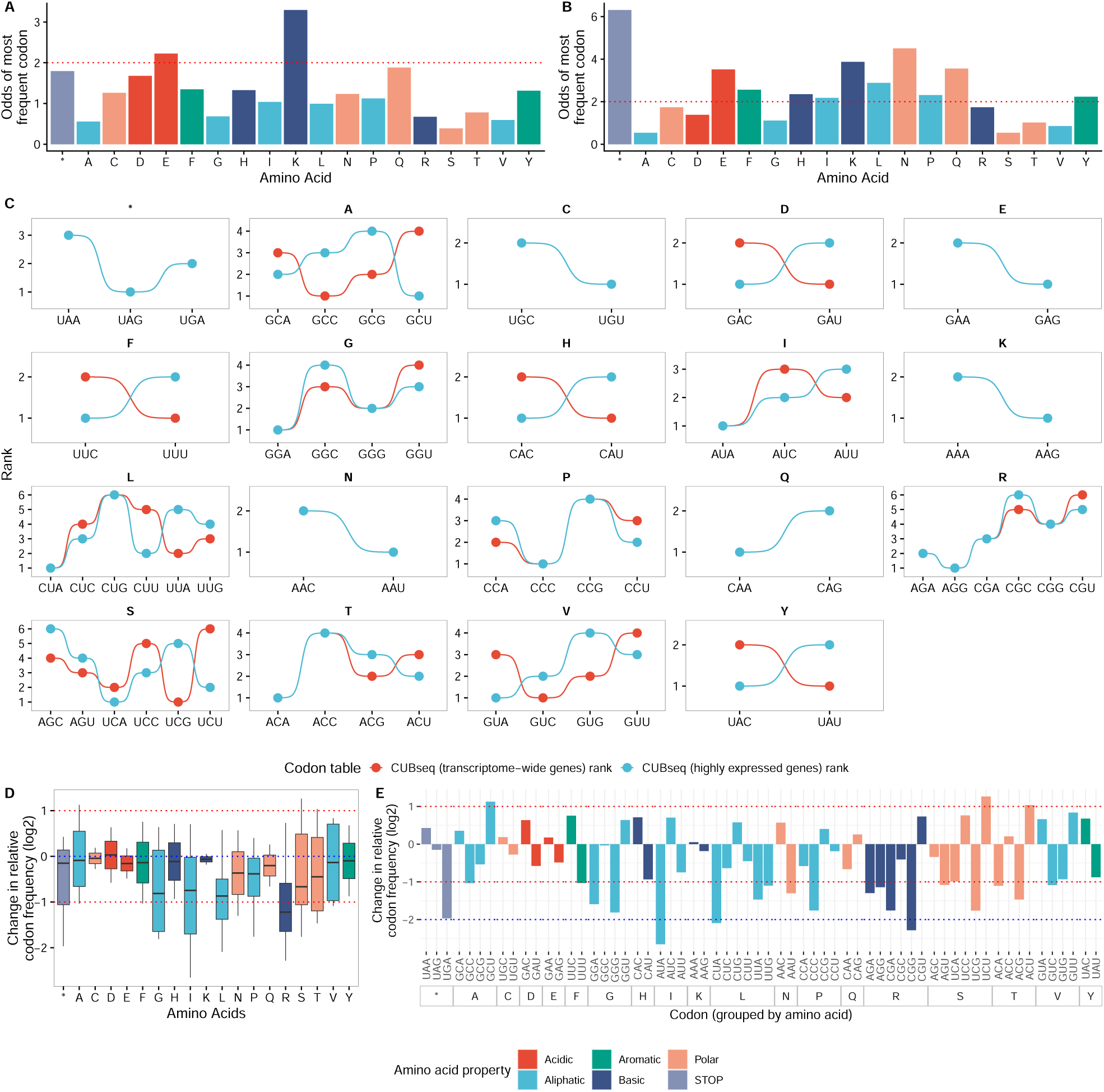
Comparison of codon relative frequencies in CUBSEQ transcriptome-wide and highly expressed genes. Signature of codon preference in CUBseq **(A)** transcriptome-wide and **(B)** highly expressed genes, represented as the odds of the most frequently used codon per amino acid versus other synonymous codons. Red dotted line is used to indicate an odds threshold of 2. **(C)** Rank comparison of codon relative frequencies for each respective amino acid (and STOP codons) using all CUBseq transcriptome-wide protein-coding genes versus highly expressed genes. Rank 1 indicates codon with the lowest relative frequency. **(D)** Amino acid and **(E)** codon resolution comparison of log_2_ absolute difference in codon relative frequencies in transcriptome-wide versus highly expressed genes. In the latter two plots, a positive log_2_ value indicates a higher frequency of the respective codon in the CUBseq highly expressed panel of genes, while a negative log_2_ value indicates a higher codon frequency in the CUBseq transcriptome-wide Figure 6: (Continued.) panel of genes. A log_2_ value of 0 indicates no change in codon frequency. Red and blue dotted lines are used to indicate a one- and two-fold difference, respectively. For all plots Methionine and Tryptophan codons are removed.

In contrast, multiple amino acids were found to have strong codon preferences within the set of highly expressed genes, with overall preference averaging at 2.38 across amino acids (Fig. 6B). Here, we observed Asparagine to have the strongest codon preference ratio of 4.5 : 1 in favour of its *AAC* codon over its *AAU* codon. This is followed by Lysine having a preference of 3.9 for its *AAA* codon, Glutamine with a preference of 3.6 for its *CAG* codon, and Glutamate with a preference of 3.5 for its *GAA* codon. Leucine, Phenylalanine, Histidine, Proline, Tyrosine and Isoleucine also exhibit strong codon preferences greater than 2 for their most frequent codons. Similar to transcriptome-wide genes, Serine, followed by Alanine and Valine displayed the lowest preference for their most frequent codon, *UCU* (0.54), *GCU* (0.54) and *GUU* (0.85) respectively.

These findings indicate that the selective pressure on codon usage exhibits diverse strengths and specificities among different amino acids, operating at both the transcriptome-wide level and within highly expressed genes. Importantly, we find low variance in codon preference within transcriptome-wide genes and greater variance within highly expressed genes, suggesting greater codon bias and stronger selective pressures in highly expressed genes, even when considering multiple experimental conditions.

We also found substantial differences in estimated codon relative frequencies when comparing our CUBseq HEG panel and CUBseq transcriptome-wide protein-coding genes. For example, we found major differences when comparing the relative ranks of codon frequencies for each amino acid (Fig. 6C). In particular, complete agreement in codon usage ranks were observed for STOP codons, as well as two-codon amino acids: Cysteine, Glutamate, Lysine, Asparagine and Glutamine. We observed a similar trend in amino acids such as Glycine, Isoleucine, Proline, Arginine and Threonine. However, several amino acids showed major differences in codon relative frequency ranks, such as Aspartate, Phenylalanine, Histidine and Tyrosine (two-codon); Alanine and Valine (four-codon); and Leucine and Serine (six-codon).

We then sought to determine the extent of variation within codon frequencies at the amino acid level and determine which codons were driving these observed changes. At the amino acid level, we found major differences when comparing codon relative frequencies within CUBseq transcriptome-wide genes versus HEG (Fig. 6D), with Arginine showing the most variation at just above one-fold difference. Amino acids Aspartate, Alanine, Cysteine and Lysine showed negligible average difference. At the codon level, we find that for some amino acids multiple codons are driving the observed differences in codon frequencies (for example, Arginine, Glycine, Lysine, Alanine), whereas for some amino acids single specific codons are driving these differences such as codon AUA encoding amino acid Isoleucine (for brevity, we will express this as Isoleucine*_AUA_*), Leucine*_CUA_*, and Proline*_CCC_* (Fig. 6E). We found that Alanine*_GCU_*, Serine*_UCU_* and Threonine*_ACU_*, each had a greater than one-fold difference in usage within the CUBseq HEG panel. In total, 20 codons displayed greater than one-fold difference, 3 of which displayed over a two-fold difference, with majority showing more limited differences compared with the CUBseq transcriptome-wide panel of genes. Furthermore, despite amino acids (and STOP codons) having complete agreement in rankings of codon usage preference across CUBseq transcriptome-wide and highly expressed genes, we still observed significant differences in usage. For example, STOP*_UGA_* is used almost two-fold more in transcriptome-wide genes than in HEG.

Our results suggest that highly expressed genes have significantly different codon usage frequencies compared to the transcriptome-wide estimates, as well as significantly greater codon usage bias. This also has implications for applications in biotechnology, whereby gene expression sustained at a high level might be achieved by recoding genes to mimic the codon usage (excluding ribosomal genes), rather than the species-level transcriptome-wide estimates.

### Transcriptome-wide codon usage shows limited strain-to-strain variability

CUBseq employs a per-sample sequencing approach, allowing us to construct an aggregated view of the *E. coli* transcriptome that captures mutational variability across multiple strains and experimental conditions. We therefore wanted to determine to what extent our species-level CUBseq transcriptome-wide codon frequencies differed with transcriptome-wide estimates obtained from genome reference assemblies.

To do this, we calculated transcriptome-wide codon usage relative frequencies using all protein-coding genes in our 6, 763 reconstructed transcriptomes, allowing us to characterise codon usage bias at the species level, regardless of gene expression level or function. We compared our CUBSeq estimates to those obtained by CoCoPUTs for *E. coli* K-12, which analogously computes codon usage across all protein-coding genes, and found complete consensus in ranked codon preferences across all amino acids (Fig. 7A). This was also confirmed when comparing codon relative frequencies at amino acid and codon resolution, where we found negligible differences between CUBseq and CoCoPUTs transcriptome-wide estimates (Fig. 7B-C).

**Figure 7:**
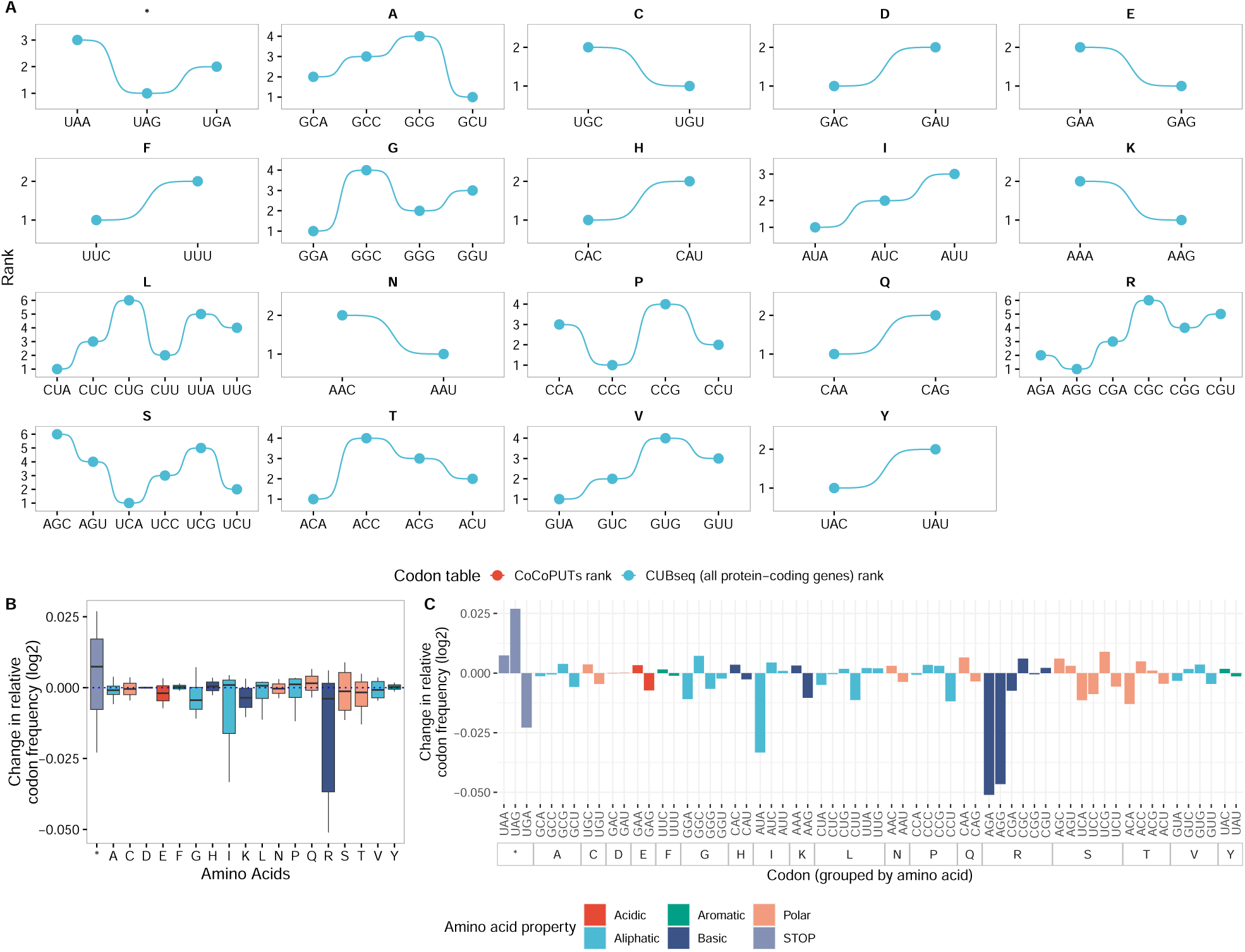
Transcriptome-wide comparison of CUBSEQ and CoCoPUTs codon relative frequencies using all protein-coding genes. **(A)** Rank comparison of transcriptome-wide codon relative frequencies for each respective amino acid (and STOP codons) in CUBseq versus CoCoPUTs. CUB-seq, employing a per-sample sequencing approach, represents an aggregated transcriptome encompassing multiple samples. Whereas CoCoPUTs utilizes a single genome reference for its analysis. Rank 1 indicates the codon with the lowest relative frequency. **(B)** Amino acid and **(C)** codon resolution comparison of log_2_ absolute difference in transcriptome-wide CUBseq versus CoCoPUTs codon relative frequencies. In the latter two plots, a positive log_2_ value indicates a higher frequency of the respective codon in the CUBseq transcriptome-wide panel of genes, while a negative log_2_ value indicates a higher codon frequency in the CoCoPUTs panel of genes. A log_2_ value of 0 indicates no change in codon frequency. For all plots Methionine and Tryptophan codons are removed.

Taken together, our results suggest limited strain-to-strain variability in codon usage, indicating a consistent transcriptome-wide signature at the species level and in the absence of strong selective pressures. Furthermore, genomic mutations have limited impact on codon usage estimates when considering all protein coding genes, as they are usually a negligible fraction of the whole transcriptome.

### CUBseq highly expressed genes show major differences in codon frequencies compared to alternative codon tables

To compare how codon usage bias differs in our CUBseq panel of highly expressed genes with alternative codon usage tables, we compared our CUBseq codon relative frequencies with that of Kazusa and CoCoPUTs *E. coli* K-12 codon relative frequencies. Both Kazusa and CoCoPUTs represent codon tables that are non-selective for the gene panel, with Kazusa being based on 14 CDS and CoCoPUTs being based on all transcriptome-wide protein coding genes. Furthermore, unlike CUBseq, neither Kazusa or CUBseq take a per-sample approach but instead are based on a single genome assembly.

At the amino acid level, we found major differences when comparing codon relative frequencies. Rank comparison of codon frequencies across the CUBseq HEG panel, Kazusa and CoCoPUTs codon usage tables showed certain amino acids exhibiting more variation than others (Fig. 8A). Amino acids (with two codons) found to show complete agreement in ranks across tables were Cysteine, Glutamate, Lysine, Asparagine and Glutamine, along with all STOP codons. Some amino acids (with *>* 2 codons) showed general agreement in ranks, such as Glycine, Proline, Arginine and Threonine. In contrast, amino acids Alanine, Aspartate, Phenylalanine, Histidine, Isoleucine, Valine and Tyrosine had matching ranks across Kazusa and CoCoPUTs tables, but differed nearly completely when compared to CUBseq HEG codon frequencies. Finally, amino acids Leucine and Serine showed highly disagreeable ranks.

**Figure 8:**
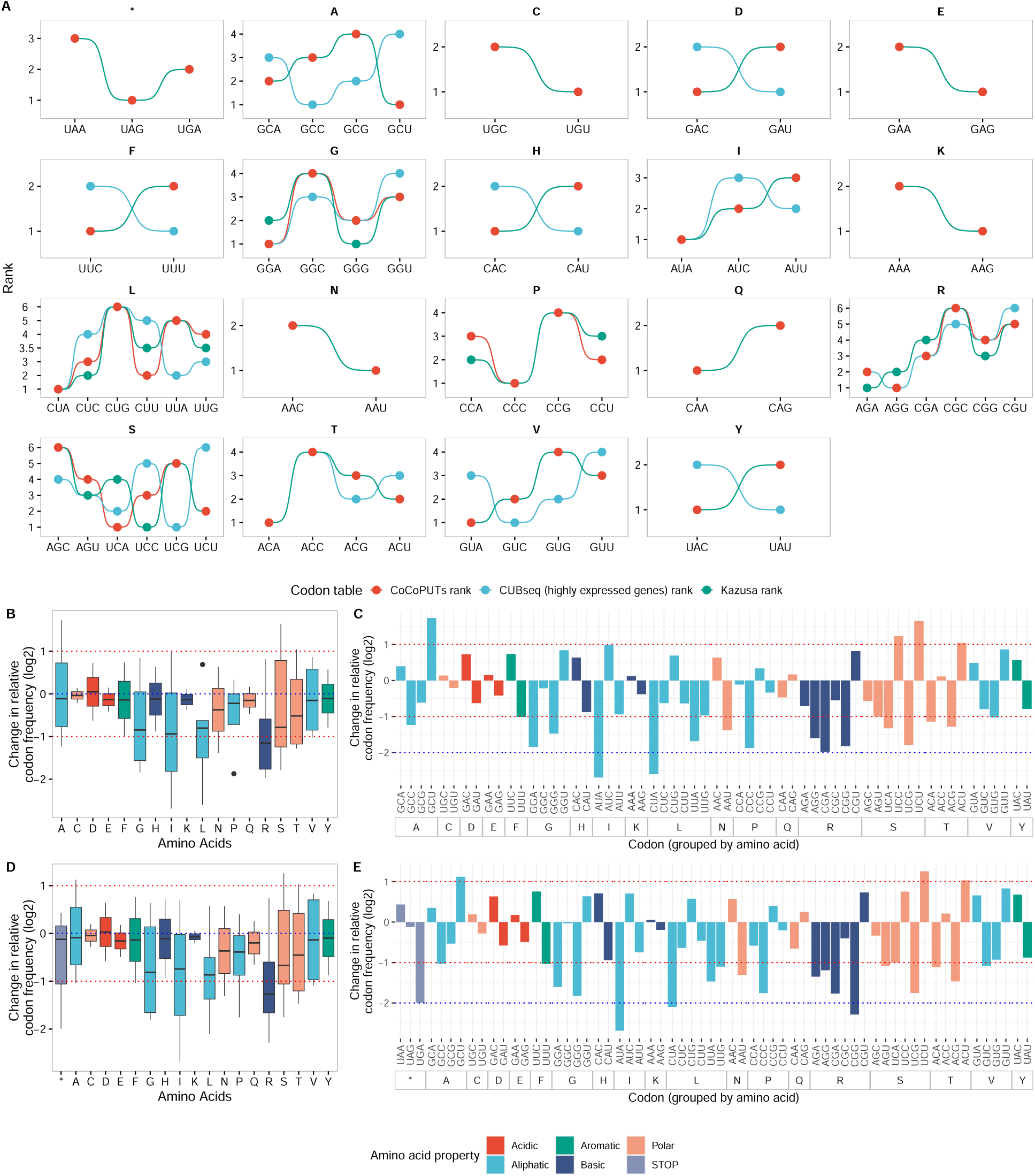
Comparison of *E. coli* K-12 relative codon frequencies in CUBSEQ highly expressed genes versus Kazusa and CoCoPUTs genes. **(A)** Rank comparison of codon relative frequencies for each respective amino acid (and STOP codons) across each codon usage table: CUBseq highly expressed genes (shown blue), CoCoPUTs (red) and Kazusa (green). Rank 1 indicates the codon with the lowest relative frequency. **(B)** Boxplots comparing distribution of log_2_ absolute differences in relative codon frequency between CUBSEQ and Kazusa and **(D)** CoCoPUTs genes. Each box represents the log_2_ absolute difference in relative codon frequencies at the amino acid level. Blue and red dotted lines indicate a zero- and one-fold difference, respectively. Boxplots show the median (thick centre line), along with interquartile ranges (box, 25% to 75%) and whiskers extending 1.5x the interquartile range. **(C)** Comparison of codon relative frequencies in CUBSEQ highly expressed genes versus Kazusa and **(E)** CoCoPUTs genes at codon resolution. Each bar represents the log_2_ absolute difference in codon relative frequency, grouped by amino acids. Red and blue dotted lines are used to indicate a one- and two-fold difference, respectively. For the latter four plots, amino acids and codons are coloured by their corresponding amino acid property. Methionine and Tryptophan codons are removed for all plots; STOP codons are removed for comparison with Kazusa as codon “UAG” is reported as zero. A positive log_2_ value indicates a higher frequency of the respective codon in the CUBseq highly expressed panel of genes, while a negative log_2_ value indicates a higher codon frequency in the Kazusa and CoCoPUTs panel of genes, respectively. A log_2_ value of 0 indicates no change in codon frequency between the two codon usage tables.

Closer inspection of how codon relative frequencies compared at the amino acid level revealed significant differences in frequencies for several amino acids. Most notably Arginine, showed a greater than one-fold difference across both Kazusa (Fig. 8B) and CoCoPUTs (Fig. 8D) codon tables relative to CUBseq HEG, followed by Isoleucine, Glycine, Leucine and Serine. Whereas Cysteine, Glutamate, Lysine and Glutamine showed negligible variation across both codon tables.

At the codon level, we found differences in frequencies being driven by individual or multiple codons for each amino acid, with similar patterns emerging across both Kazusa (Fig. 8C) and CoCoPUTs (Fig. 8E) codon tables relative to CUBseq HEG. For example, we find that codon Isoleucine*_AUA_* was found to be the main driver of differences observed for this amino acid, exhibiting the largest change in frequency, −2.69 and −2.68 in Kazusa and CoCoPUTs, respectively. Leucine*_CUA_* (Kazusa: −2.60, CoCoPUTs: −2.10) and Arginine*_CGA_* (Kazusa: −1.98, CoCoPUTs: −2.28) were also found to act as the main drivers of variation observed between codon usage estimates. In contrast, amino acids such as Arginine and Serine have multiple codons contributing to relative codon frequency differences observed for their respective amino acids. Furthermore, greater variation is observed for the Arginine*_CGA_* codon in the Kazusa table, whereas Arginine*_CGG_* is driving the variation in the CoCoPUTs table (reaching greater than two-fold difference).

In summary, our transcriptome-derived codon relative frequencies based on our panel of CUBseq HEGs display consistent patterns in their relative codon usage frequencies when compared with alternative single-reference-based codon usage tables such as Kazusa and CoCoPUTs. In particular, when comparing overall relative codon frequencies at amino acid resolution, we found Arginine, Isoleucine, Leucine and Glycine seem to exhibit the most variation in relative codon frequencies. At codon resolution, we find specific codons driving these codon frequency differences for some amino acids. Such differences could be exploited for biotechnological applications such as codon optimisation for heterologous protein expression, and target amino acids that may require greater fine-tuning.

### Codon relative frequencies often correlate with number of cognate tRNAs

To demonstrate the potential applications of our tool in analysing codon usage at the transcriptome level, we explored the relationship between codon relative frequencies of transcriptome-derived HEGs and their cognate tRNA gene abundances. Using our tRNA gene expression data to infer tRNA abundance levels, and tRNA gene names and codon pairings obtained from the GtRNAdb, we calculated a TPM rank score to represent tRNA gene abundance. We then mapped each tRNA abundance score rank to its corresponding codon relative frequency rank (with rank 1 indicating the greatest tRNA gene abundance and codon relative frequency, respectively) (Fig. 9).

**Figure 9:**
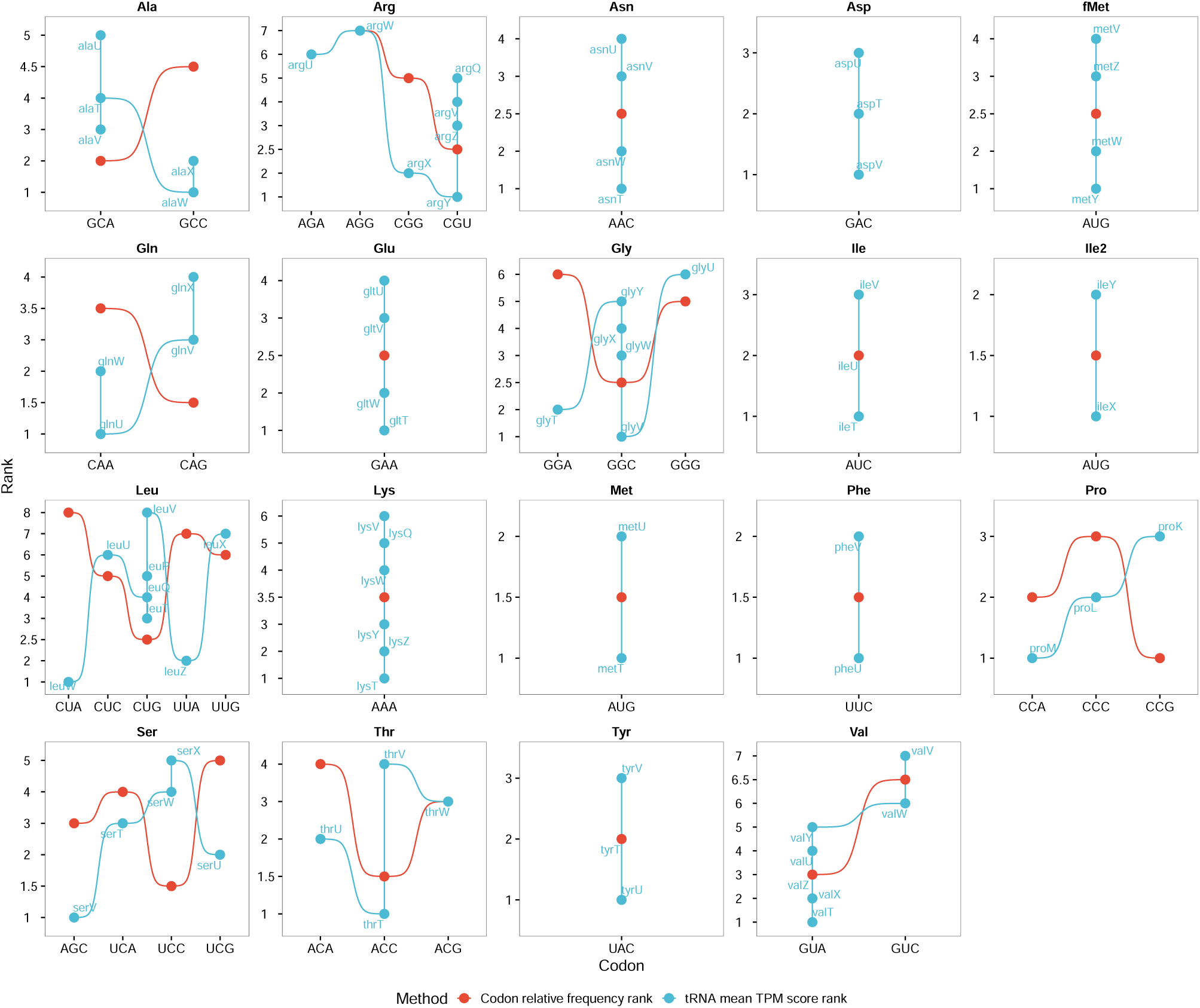
Relationship of tRNA gene abundance with corresponding codon relative frequencies of transcriptome-derived highly expressed genes. Each panel represents an amino acid with tRNA gene abundance mean score (shown in blue and labelled by tRNA gene) and the respective codon relative frequency (shown in red). For example, the amino acid Glutamine (”Q”) has two codons *CAA* and *CAG*, which are recognised by tRNAs *glnW*, *glnU* and *glnX*, *glnV*, respectively. The panel therefore shows tRNA gene abundance score rank compared to its corresponding codon relative frequency rank. Rank 1 indicates the highest tRNA gene abundance score and highest codon relative frequency, respectively. Here, tRNA gene abundance rank (blue) is compared to its respective codon relative frequency rank (red) in transcriptome-derived CUBSEQ panel of significantly highly expressed genes. Therefore, matching ranks indicate a positive relationship between tRNA gene abundance and its corresponding codon relative frequency. Note that some codons are recognised by more than one cognate tRNA gene, leading to some codons containing more than one tRNA gene with the same anticodon (see Glutamine example above).

We found general agreement in ranks, or positive relationships between codon relative frequency and tRNA abundance mean scores within the tRNA-Arginine, tRNA-Threonine and tRNA-Valine gene families. Intriguingly, we found that codons exhibiting the highest relative frequency for their respective amino acid were generally frequently associated with a larger repertoire of tRNAs, with the exception of Proline codons. For example, within the tRNA-Arginine gene family, codon CGU is recognised by four different cognate tRNA genes, with *argY* having the greatest tRNA abundance, followed by *argZ*, *argV* and *argQ*. The rest of the Arginine codons are recognised by one cognate tRNA (not accounting for wobble).

Opposite associations of ranks, or negative relationships were observed in the tRNA-Alanine, tRNA-Leucine, tRNA-Proline, tRNA-Glutamine and tRNA-Serine gene families. Despite this, we observe similar patterns with codon frequency and tRNA abundance. For example in Serine, we observe codon UCC to have the highest relative frequency and is recognised by two tRNA genes, with *serW* displaying greater abundance, followed by *serX*.

Although we cannot determine the causal direction between codon relative frequency and tRNA gene abundance, these findings warrant further analysis into the potential mechanisms driving the relationship between codon usage and tRNA recognition.

## Discussion

In this work we developed CUBSEQ, a pipeline that performs large-scale transcriptome-wide codon usage analysis by processing RNA-sequencing reads using a unified and standardized workflow, enabling the comparison of hundreds of studies across different experimental conditions and in any species where RNA-sequencing data is available (Fig. 1). Using this approach we performed, to our knowledge, the first and most comprehensive transcriptome-wide codon usage analysis on *E. coli* by incorporating per-strain and per-sample variance, allowing us to construct a composite transcriptome to account for this sequence variation. By leveraging the rapid growth of transcriptomic data, we overcome a common limitation of previous codon usage studies, which rely on a single reference genome to derive codon frequency estimates. Our variant analysis allows us to account for the presence of mutations in the transcriptome, which we found to be widespread in *E. coli* (Fig. 2A), with a significant proportion of genes found to be recurrently mutated relative to the *E. coli* K-12 reference genome (Fig. 2B). Therefore, the consistent presence of mutations across the transcriptome argues a strong case towards adjusting codon usage estimates to account for the inherent sequence variation observed within genes across thousands of samples.

Current codon usage analysis methods are often non-selective on the panel of genes being studied. To address this challenge, we developed a novel meta-analysis approach to identify a panel of genes that are consistently highly expressed across sample library size, experimental conditions and robust to strain variation (Fig. 3B). We find that a majority of these genes are significantly enriched for biological processes such as metabolism and translation (Fig. 3A), giving us further confidence in our selected panel of genes. Our approach of identifying and adjusting codon usage estimates towards highly expressed genes, whilst capturing the sequence variations within them, represents a valuable tool for driving advancements in biotechnology, as our method lays the foundation for tailored strategies to enhance heterologous protein expression by identifying and aligning codon usage estimates towards genes with robust and consistent expression patterns at the transcriptional level.

We find major differences when comparing transcriptome-wide frequencies with our identified panel of HEGs (Fig. 6), demonstrating the large effect the panel of genes can have on codon usage estimates and with greater codon usage bias observed in HEGs having implications for codon optimisation. This contrasts with the negligible differences observed between CUBseq and CoCoPUTs when considering all transcriptome-wide protein-coding genes (Fig. 7), suggesting that codon usage in *E. coli* is consistent across strains and experimental conditions and when using transcriptome-wide protein coding genes.

Furthermore, CUBSEQ has demonstrated that there are major differences in estimated codon frequencies derived from CUBseq HEG versus alternative codon tables (Kazusa and CoCoPUTs) that are non-selective in their gene panels and are based on a single genome assembly. In particular, Arginine codon frequencies showed the most deviation. However, at the codon level we see individual or several codons may be the drivers of large variations observed at the amino acid level (Fig. 8). It is also interesting to see consistent patterns emerging at amino acid and codon resolution across the Kazusa and CoCoPUTs tables, despite both using entirely different gene sets, which could suggest a conserved “baseline” codon usage signature that exists when selection is weak.

We then demonstrate one of the many potential broad applications of transcriptome-based codon usage estimates, and assess the relationship with their cognate tRNA gene abundance. Here, we find codon relative frequencies often correlating with their cognate tRNA abundances. However, we also find many negative correlations between codon frequencies and their cognate tRNA gene abundance. For codons recognised by more than one tRNA gene, we also identified differing tRNA gene abundance patterns despite recognition of the same codons (Fig. 9), and a general trend where high frequency codons were often associated with a larger reportoire of tRNA recognition. These results could be extended by analysing codon autocorrelation patterns to explore their association with tRNA recycling (as outlined in [14]). It is important to note that this study does not account for wobble and near-cognate codons in its analysis, so some codons are not assigned a tRNA, however this should be explored in future studies.

The main challenge of this study is the accurate capture and quantification of codons and minimising sources of codon skew. In calculating relative codon frequencies, we assume all genes to be consistently up-regulated as would be in wild-type, however as with any meta-analyses, it is difficult to manually filter out studies with engineered strains (for example, studies that have performed gene knockout and mutation experiments). Although, it could also be argued that due to the large sample size (6, 763 sequencing runs), these engineered strains are far outweighed by wild-type, and so may not make a difference in calculated relative frequencies.

The potential for further analysis is vast. Although this study only looks at protein-coding genes, assessing codon usage profiles across regulatory RNA elements could uncover further mechanisms towards their role in gene regulation and protein expression. As the central crux of this study is on the influence of synonymous variants on codon usage, it would also be interesting to decode mutational synonymous codon-switching events (as inspired by [23]) at the transcriptome-wide level as well as based on gene panels, in comparison to our composite reference transcriptome. Furthermore, in a similar spirit to Novoa et al, who identified that Arginine is able to separate species into their corresponding domains [13], it would be interesting to determine whether transcriptome-derived codon usage can be used to identify strain-level codon usage signatures. Studies have also increasingly explored the effects of synonymous codon usage on protein structure, with one recent study examining more local associations on the distribution of amino acid backbone dihedral angles [16]. It would therefore be interesting to see what insights transcriptome-based codon usage estimates can give to protein structure dynamics.

One particularly important extension of this study would be to perform codon “optimisation” or gene recoding experiments, using the CUBseq-derived codon frequencies shown for all protein-coding genes (Fig. 4) and HEGs (Fig. 5). As well as testing the effects on protein expression and solubility, as CUBseq codon usage estimates are derived at the transcriptional level, it would be interesting to observe the effects of CUBseq-recoded genes on any translation-dependent and translation-independent cell functions. mRNA stability would also be a good candidate feature to test for, based on their AT3 and GC3 content - where we would expect HEGs to be more GC3-rich, as it has been shown to proportionately increase GC content and confer mRNA stability [24].

Overall, CUBSEQ provides a reproducible and high-throughput addition for performing large-scale codon usage analysis at the transcriptome level. Accurate quantification of codon usage is essential to applications in codon optimisation to increase protein yield in heterologous protein expression, as well as studies in transcriptional and translational regulation, in diseases such as cancer, and exploring the molecular evolution of species and host/pathogen relationships. The incorporation of up-to-date transcriptome abundance information to infer gene expression makes CUBSEQ suitable for a wide-range of codon usage analyses.

## Methods

### Data retrieval, curation and pre-processing

RNA sequencing data was retrieved from the European Nucleotide Archive (ENA) through its REST API [25]. Specifically, we considered all paired-end sequencing runs associated with the root taxa ID for *E. coli* (562) and considered only experiments annotated with TRANSCRIPTOMIC as library source, ILLUMINA as instrument platform and RNA-Seq as library strategy. We further restricted our analysis to data generated from 2012-01-01 to 2022-05-31, ultimately leading to an initial dataset of 6, 983 sequencing runs.

The initial dataset was further filtered by removing samples with single FastQ files, which likely denote interleaved paired-end or mislabelled single-end reads. Samples with raw data more than 55GB in size were also discarded, as inferred from the sra bytes field, to limit the computational burden of both the alignment and quantification steps. A further 214 sequencing runs were discarded due to various technical issues during processing (see additional files for removed sequencing runs along with justification). In total, 220 sequencing runs were removed (*≈* 3.15% of the initial dataset), leaving a total of 6,763 sequencing runs considered for downstream analysis (see Supplementary for additional details on filtering criteria as well as the metadata file containing all study accession numbers used in this study).

Finally, paired-end reads in the final dataset were pre-processed with fastp [26], using default parameters to remove adapters and low quality bases.

### Transcript-level quantification

We used Salmon [27] (v1.9.0) to perform transcript level quantification using the pre-processed raw reads as input with automatic library type detection, and a reference transcriptome generated from the wild-type reference sequence and corresponding annotation using the gffread package [28].

We then summarised the Salmon quantification results at the gene level using the tximport package [29], while setting the --counts-from-abundance parameter to no and leaving all other parameters to their default values. Ultimately, this generates a count matrix, where we extracted genes and their estimated abundance expressed as Transcripts Per Million (TPM).

### Transcriptome reconstruction

We used the *E. coli* str. K-12 substr. MG1655 genome and GTF annotation files (Ensembl, version ASM584v2, download date: 27*/*04*/*2022) as the reference genome for our analysis. Pre-processed reads were aligned to the reference *E. coli* genome using the splice-aware aligner STAR (v2.7.10) [30]. To do this, we first built a genome index by setting genomeSAindexNbases to 10, which was determined using the recommended scaling formula and setting GenomeLength to 4.6*M* bp. We also set alignIntronMax to 1 to disable spliced alignment of unannotated junctions. All other parameters were set to their default values.

The generated alignment was then used to identify single nucleotide variants (SNVs) and indels using the bayesian haplotype-based variant detection tool freebayes [31], by setting the --ploidy parameter to 1 while keeping all other parameters to the corresponding defaults. We then used the bcftools [32] norm command to normalize variants, decompose multiple nucleotide variants into consecutive SNVs, and retain those with a Phred score quality above 20. To account for indels causing a frameshift in the sequence (thus resulting in completely different sequences of codons/amino acids and skewing codon counts) variants were filtered to retain only SNVs.

Finally, the filtered set of variants were used to generate a sample-level transcriptome using the consensus command in bcftools.

### Mutational analysis

We performed variant quality control by merging VCF files from each sample using bcftools merge. To prevent memory overload when loading the merged VCF file, genotypes were removed prior to analysis using the bcftools view command with the --drop-genotypes option. We then parsed the resulting merged VCF file using the VCFR package [33], and filtered manually to compute statistics across the *E. coli* genome on alternate allele observation count (AO), read depth (DP), mean mapping quality of observed alternate alleles (MQM), Phred-scaled quality (QUAL) and the average number of mutations per sample across each individual loci (AC).

To account for the large distribution of AO counts and facilitate data interpretability, AO counts were set to a threshold at the 99.9th percentile of the distribution to avoid skew by extreme AO values. Similarly, DP and QUAL values were log_10_-scaled. To determine the average number of mutations, the AC column (representing, for a given locus, the total number of alternate alleles in called genotypes) was normalised by dividing by the total number of sequencing runs.

It is important to note, for the purposes of this study, we filter out non-synonymous mutations to avoid frameshift mutations and their consequential skewing of codon counts. However, the resulting mutations cannot be considered true “synonymous” mutations due to relying on a single reference genome. It is therefore more accurate to consider the alternate alleles identified in our analysis as “likely synonymous” relative to our reference genome, but still likely incorporating non-synonymous mutations as well.

To determine the total number of mutations per gene across sequencing runs, the filtered VCF files for each sample were further annotated with bedtools annotate by passing the GTF file and using the --both flag to provide both the counts and fraction of counts. Finally, we consolidated this information to get the number of samples carrying a mutation in a given gene.

### Determining transcriptome-wide genes

For transcriptome-wide codon usage analysis, we followed the same/similar filtering approach as CoCoPUTs where we retained all protein-coding genes and filtered to retain “CDS” with ribosomal genes filtered out (i.e. ribosomal subunit proteins). This was achieved by first identifying all ribosomal proteins from the GFF file (assembly version: ASM584v2). This was done by filtering the “type” field for “gene”, “biotype” for “protein coding”, and filtering the “description” field to keep any proteins containing the strings “ribosomal subunit protein” and “putative ribosomal protein”. The remaining ribosomal-associated enzymes (although not ribosomal proteins) were removed by filtering any strings containing “ase” in the “description” field, giving us a total of 57 ribosomal proteins (including 2 putative ribosomal proteins) that were removed. In addition, a further 5 isoforms and 39 transposable elements were removed. Genes were also checked to have a CDS divisible by 3. This gave us a final set of 4, 141 protein-coding genes, the IDs of which were used to subset the TPM gene expression matrix. The relative codon frequencies were subsequently calculated (as outlined below) and used for downstream analysis.

### Determining highly expressed genes

To determine a panel of highly expressed genes, we used a non-parametric meta-analysis approach. We first filtered each sample to retain only the non-ribosomal protein-coding genes (as described above) and subset the TPM matrix to keep only these genes. For each sample, we then computed a gene score as the rank of their TPM value, such that genes with a higher TPM have a higher rank, which is in turn transformed into z-scores using the inverse normal transform; this allows us to determine consistently highly expressed genes regardless of differences in experimental conditions, library sizes across samples and whether the distribution of gene expression levels are not normal. We denote as consistently highly expressed those genes with an average score *≥* 1.644, which corresponds to the upper 95% quantile of the normal distribution.

To compute the statistical significance of the scores associated with our high expression genes, we converted per sample gene scores to *χ*^2^ values with 1 degree of freedom, which were then summed to obtain a new score that is *χ*^2^ with *n* degrees of freedom, with *n* being the number of samples. The resulting p-values were then adjusted for multiple hypotheses testing using the Benjamini-Hochberg procedure, and denoting significant genes with FDR *<* 0.01.

Using this approach, we also identified significant low expression genes, however this carries many caveats in the context of RNA-sequencing. For example, genes observed to be lowly expressed could be affected by batch effects, be a result of low mapping coverage, or gene length bias which particularly affect TPM counts (with longer genes tending to have more sequencing reads than shorter genes), finally small sample sizes carry inherently low statistical power making it difficult to determine whether genes are truly lowly expressed. For these reasons, we do not include low expression genes in our main analysis.

Finally, we characterised our highly expressed gene panel using Gene Ontology (GO) analysis (Released 2022-07-01) restricted to the the biological process terms [34, 35]. We used the PANTHER Overrepresentation Test (Released 20221013), with annotation from PANTHER version 17.0 (Released 2022-02-22), GO Ontology database (Released 2023-03-06) and *E. coli* as the reference list [36, 37]. To identify statistically significant GO terms, we set test type to Fisher’s Exact and a correction threshold of FDR *<* 0.05.

### Determining concordance of highly expressed genes per strain

We performed concordance at the top (CAT) analysis as outlined in [38] on significant highly expressed genes within the top taxonomic groups. We first selected the top 5 frequently occurring taxonomic groups from the metadata, and filtered our TPM gene expression matrix to only include sequencing runs from those taxonomic groups. We then performed our meta-analysis on the TPM data for each taxonomic group to identify significant highly expressed genes, with lowest ranks associated with the most significantly highly expressed gene. Using our CUBSEQ panel of significant highly expressed genes as reference, we then determined the concordance of gene rankings per taxonomic group.

### Retrieving codon usage frequencies from the Kazusa and CoCoPUTs databases

To compare CUBseq with the Kazusa [19] and CoCoPUTs [39, 20] codon usage tables, we programmatically obtained the respective codon counts for *E. coli* K-12. The Kazusa and CoCoPUTs Codon Databases contain codon counts and codon usage data (expressed in frequencies per thousand), with values calculated from GenBank (with CoCoPUTs additionally containing data from RefSeq). As we are comparing codons by their relative frequencies for this study, we only needed the raw codon counts, which were programmatically obtained from the CUTG database via FTP protocol for Kazusa, and via URL for CoCoPUTs. We filtered the respective codon count tables for taxonomy ID “83333” (*E. coli* K-12) to compare CUBSEQ codon frequencies with, and calculated relative frequency values.

### CUBseq codon usage analysis

To enable fair comparison of CUBseq with the Kazusa and CoCoPUTs codon usage tables, we used relative codon frequencies as our CUB index, allowing for comparison of codon usage across different amino acids. This can be summarised as the count of an individual codon *j* for amino acid *i*, divided by the total count of synonymous codons for that amino acid *n_i_*, so that the relative frequencies of all synonymous codons of amino acid *i* equals 1, as follows 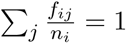.

Frequency per thousand codons were calculated by dividing the total count of a particular codon *c_i_* by the total number of all codons *C* (*C* = 128, 314, 399 codons for highly expressed genes, and *C* = 8, 968, 928, 288 for all protein-coding genes), multipled by 1000, such that: 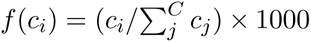.

To do this, each mutated transcriptome FASTA file (*n* = 6, 763) was filtered by transcript ID. For example, for analysis of highly expressed genes, each mutated transcriptome FASTA file was filtered to retain sequences of the 81 highly expressed genes. For transcriptome-wide analysis of all protein-coding genes, each FASTA file was filtered to retain all 4, 242 non-ribosomal protein-coding genes. Relative codon frequencies were then calculated using the codon counts generated by the Biostrings package [40] from the mutated transcriptomes of high expression genes, and all nonribosomal protein-coding genes, respectively. Finally, codon relative frequencies were then summarised to codon and amino acid levels.

To determine preference of the most frequently used codon per amino acid, we calculated the odds of the most frequent codon, such that for a given amino acid *i*, and its respective codon with the highest relative frequency (*f_ij_*_(*max*)_), we compute the likelihood of observing the most frequent codon compared to not observing it, such that: *odds* = *f_ij_*_(*max*)_*/*1 *− f_ij_*_(*max*)_.

To compare codon usage across different codon tables, we summarise codon relative frequencies at the codon and amino acid levels and calculate the absolute differences between tables.

### tRNA abundance analysis

To determine how tRNA abundance correlates with its corresponding codons, we extracted the *E. coli* K-12 MG1655 tRNAs from GtRNAdb [41, 42] (Data Release 20 [Dec 2022]) and their anticodons via URL http://gtrnadb.ucsc.edu/genomes/bacteria/Esch_coli_K_12_MG1655/ and downloaded the tRNAscan-SE results (accessed 5th March 2023). We also verified that the genome build (ASM584v2) used in this study was the same used by the GtRNAdb database (available in the “Genome Info” section).

We filtered our GTF file for tRNA genes, resulting in 86 tRNAs, and mapped the GtRNAdb tRNA anticodons (a total of 89 tRNAs) to the GTF file by their chromosomal start and end coordinates. We found, despite using the same genome assembly build, the start locus for tRNA *hisR* in GtRNAdb differed by one nucleotide to the start locus in the GTF file, and so was replaced by the start coordinate in the GTF file (3982510). The three additional tRNAs present in GtRNAdb but not in the GTF file included two “undetermined” tRNAs with “*NNN* “ as their anticodon (corresponding to isotypes *Arg* and *Ile2*, respectively), and one tRNA-Thr with anticodon CGT with a truncated start (potentially an isotype not included in the GTF). Furthermore, all three of these tRNAs had very low tRNA prediction scores and were unable to be mapped to the GTF file and thus not included in our analysis. We then converted the anticodons to their natural codons, for example, GAA (notation 5’ to 3’) would correspond to codon TTC. One limitation of this approach is that it does not consider wobble or near-cognate codon pairing, whereby there can be multiple tRNA-anticodon and codon pairings.

As gene quantification with *Salmon* was performed in alignment-free mode, the software was not able to resolve 47 of the 86 tRNA sequences due to high sequence homology. This resulted in the tRNA genes being discarded as duplicates. *featureCounts* [43] was therefore used to provide a crude gene-level quantification, by mapping the BAM files to genomic features. As CUBSEQ processes paired-end reads, we used the *-p* parameter specific to paired-end reads to count fragments instead of reads, the feature type parameter *-t* was set to count “exon” (default) in the GTF annotation file, and *-g* attribute type was set to “gene id” (default), with all other parameters being set to default values. We then derived TPM estimates from the resulting quantification files using the standard formula as denoted in [44]. As *featureCounts* does not calculate the effective length of genes, we instead used gene length to derive TPM estimates. The gene TPM quantifications were then imported to tximport and summarised at the gene level.

We then computed a normalised rank-based score of the tRNA TPMs. Here, all genes transcriptome-wide were ranked per sample and normalised by the total number of genes to generate a score. The scores were then converted into normally distributed z-scores through inverse normal transform, where the mean of the normalised scores were calculated. Genes were then filtered to only retain tRNA genes. The tRNA mean scores were then converted to ranks (with rank 1 signifying the tRNA gene with the highest mean abundance score) and plotted against their corresponding ranked codon relative frequencies (with rank 1 indicating the codon with the highest relative frequency).

### Data visualisation and development environment

All graphics were produced in R. For further details on the Nextflow workflow and development environment, see Supplementary Information.

### Data and Code Availability

All code for the CUBSEQ Nextflow pipeline and analysis are available on GitHub (also as a Docker image) and can be accessed at https://github.com/stracquadaniolab/cubseq-nf.

### Contributions

G.S. conceived the study. G.S. and A.S. designed CUBseq. A.S. implemented CUBseq and performed all experiments under G.S. supervision. G.S., A.S., J.P., and C.L. analysed the data. A.S. and G.S. wrote the manuscript with contributions from all the authors.

## Acknowledgments

This work was supported by the UKRI EPSRC Fellowship (EP/V033794/1) to G.S, and supported by the UKRI Biotechnology and Biological Sciences Research Council (BBSRC) grant number BB/T00875X/1.

