## Supplementary Materials for "Transcriptome-wide meta-analysis of codon usage in *Escherichia coli*"

#### Methods

##### Nextflow Design

A big challenge in working with ”omics” data is managing how each of the samples are tracked, organised and processed. One powerful feature of Nextflow is its ”tuple” feature, which was used so that each of the sequencing reads could be grouped back to their respective sequencing run, experiment, sample and study accessions. Another powerful feature is Nextflow's ”error strategy”, which was used to dynamically control for RAM and CPU usage, as well as to programmatically make decisions on whether to ignore any job failures after a specified number of attempts, or whether a failed job returns a specific exit code. For example, if a particularly large sequencing run requires a large amount of RAM (i.e. during read alignment or variant calling), rather than the job failing and stopping the entire pipeline, Nextflow can dynamically increase the amount of RAM in the next attempt to process that sequencing run. This ensures efficient usage of computing resources, and to only provide higher RAM/CPU power to the few sequencing runs that require it. Similarly, if a job fails and returns a specific exit code (e.g. during Salmon quantification, some reads may be of bad quality or mislabelled entirely - leading to a zero reads mapping error), rather than halting the entire pipeline, such errors can be ”ignored” by Nextflow and the resulting sequencing runs manually removed from the metadata file. Furthermore, due to storage constraints (and to make any complications more manageable and easy to debug), the pipeline was run in yearly batches (rather than in one go), the outputs of which were all pooled together prior to downstream analysis.

##### Data retrieval, curation and pre-processing

During the data processing steps, some sequencing runs had to be completely removed from the study. Three run accessions from the year 2022 (SRR17235082, SRR17235083 and SRR17235089) were removed due to insufficient RAM to complete the *freebayes* process. Likewise, seven run accessions from the year 2020 (SRR10719777, SRR10719776, SRR10719772, SRR10719769, SRR10719779, SRR10719775 and SRR10719771) and two run accessions from 2019 (DRR089608, DRR089609), were removed due to having insufficient RAM for the alignment process.

The entire study PRJNA707564 (163 sequencing runs, from the year 2021) was removed as during the transcript quantification stage, Salmon was unable to map any reads to the transcriptome, this may be due to the reads corresponding to sRNA genes (i.e. non-coding RNA that do not encode for proteins, and which cannot be used in this analysis anyway). A further 12 run accessions from the year 2019 (SRR8521474, SRR8521466, SRR8521471, SRR8521468, SRR8521475, SRR8521470, SRR8521472, SRR8521465, SRR8521467, SRR8521469, SRR8173221 and SRR8173222), and four run accessions from 2018 (ERR2276353, ERR2276351, SRR1787591 and SRR1787590), were unable to map during Salmon quantification and had to be removed.

#### Results

### Figures

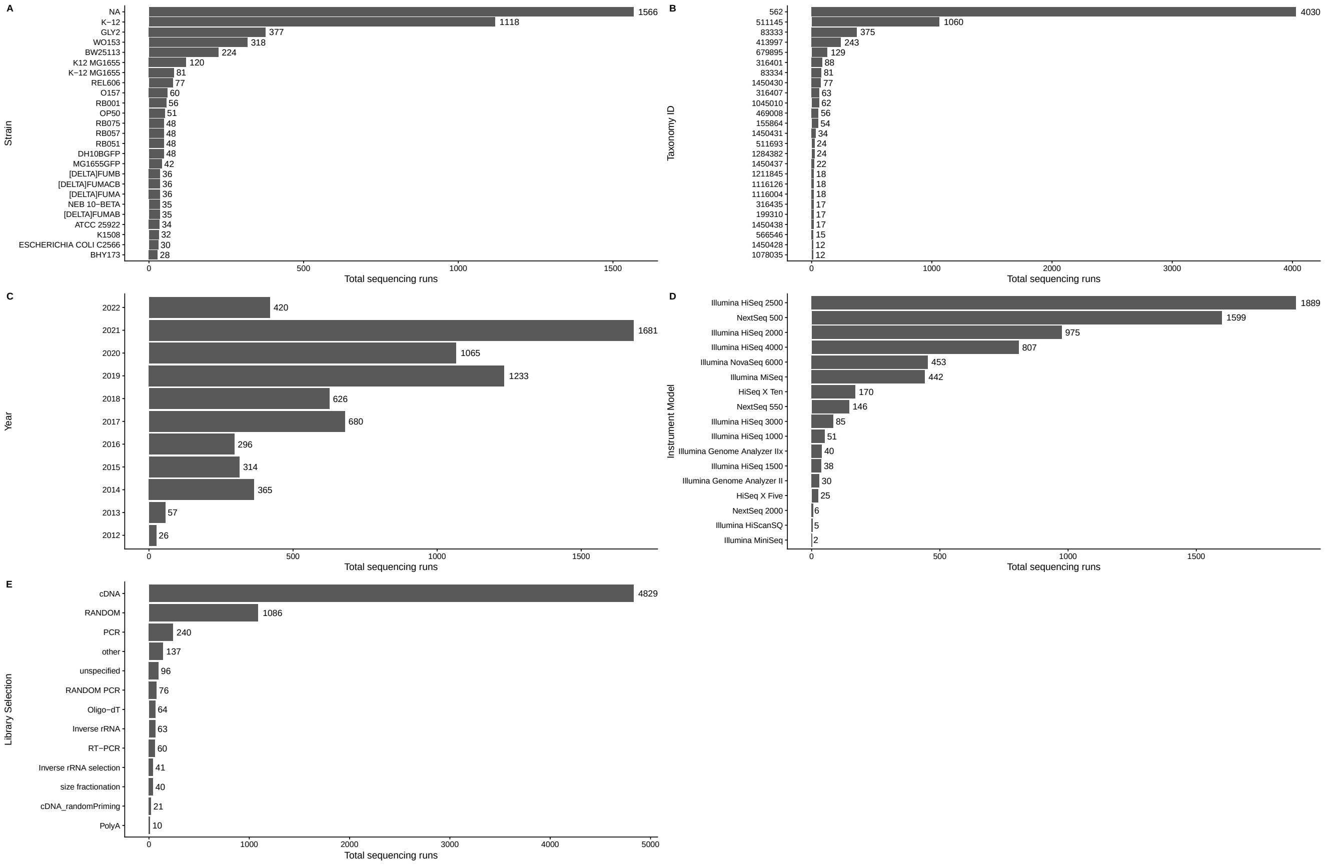


**Figure 1: Metadata descriptive statistics.**

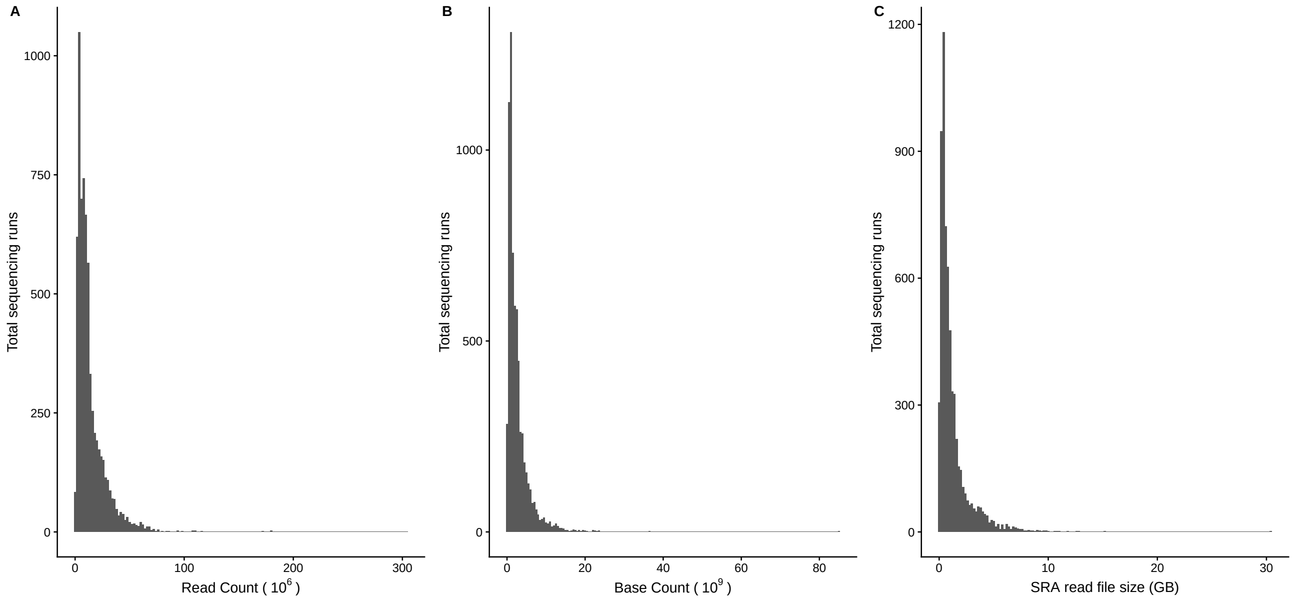


**Figure 2: Fastq statistics.**

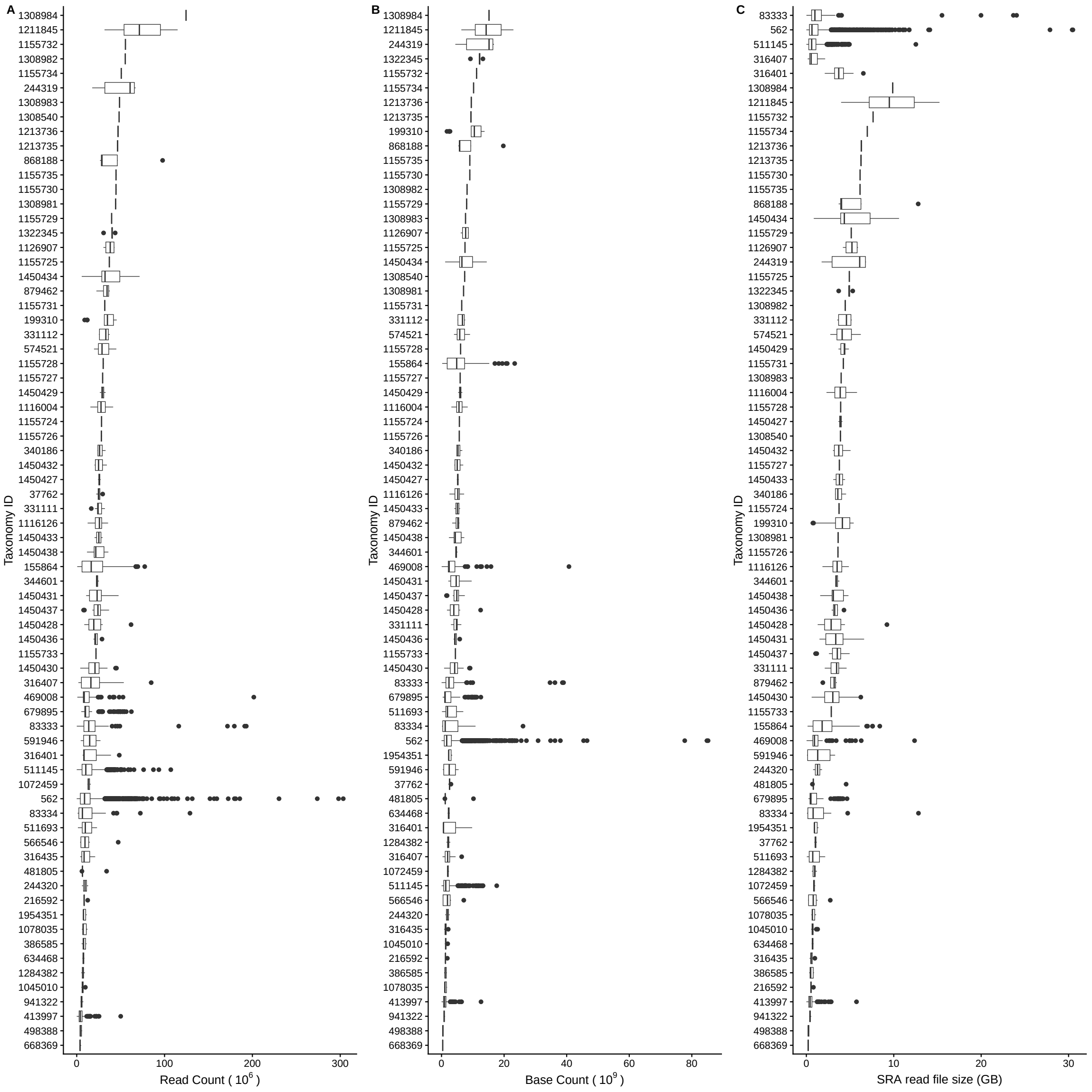


**Figure 3: Fastq statistics stratified by strain.**

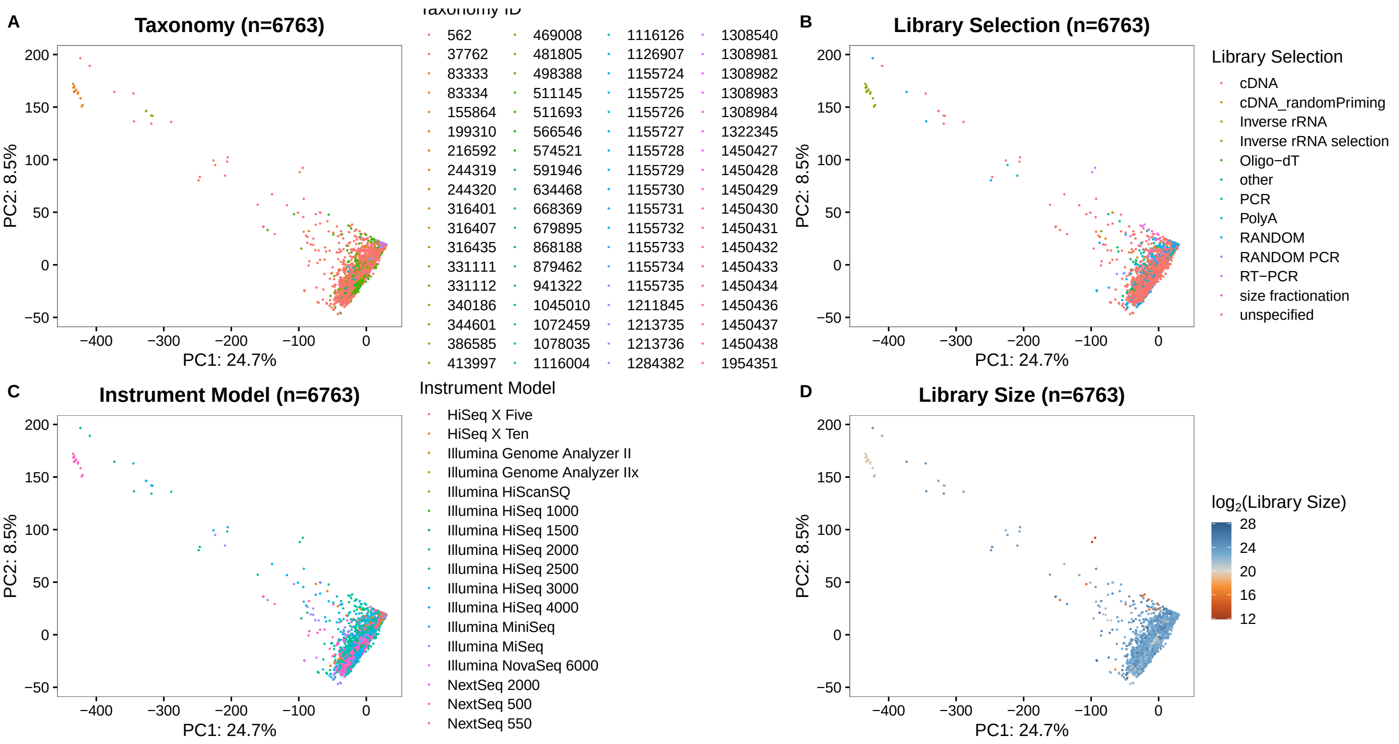


**Figure 4: PCA plot.**

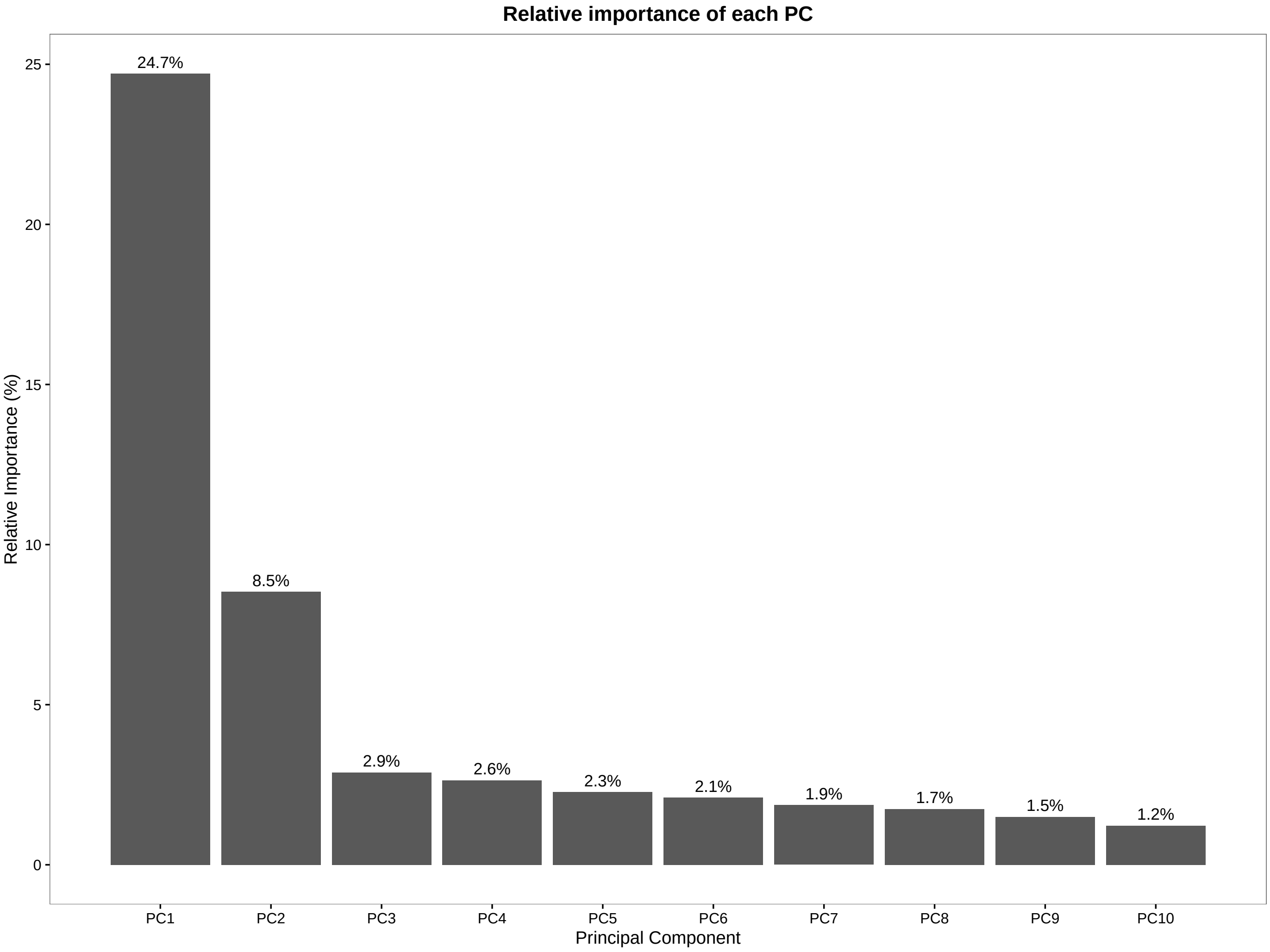


**Figure 5: Bar plot of PC relative importance.**

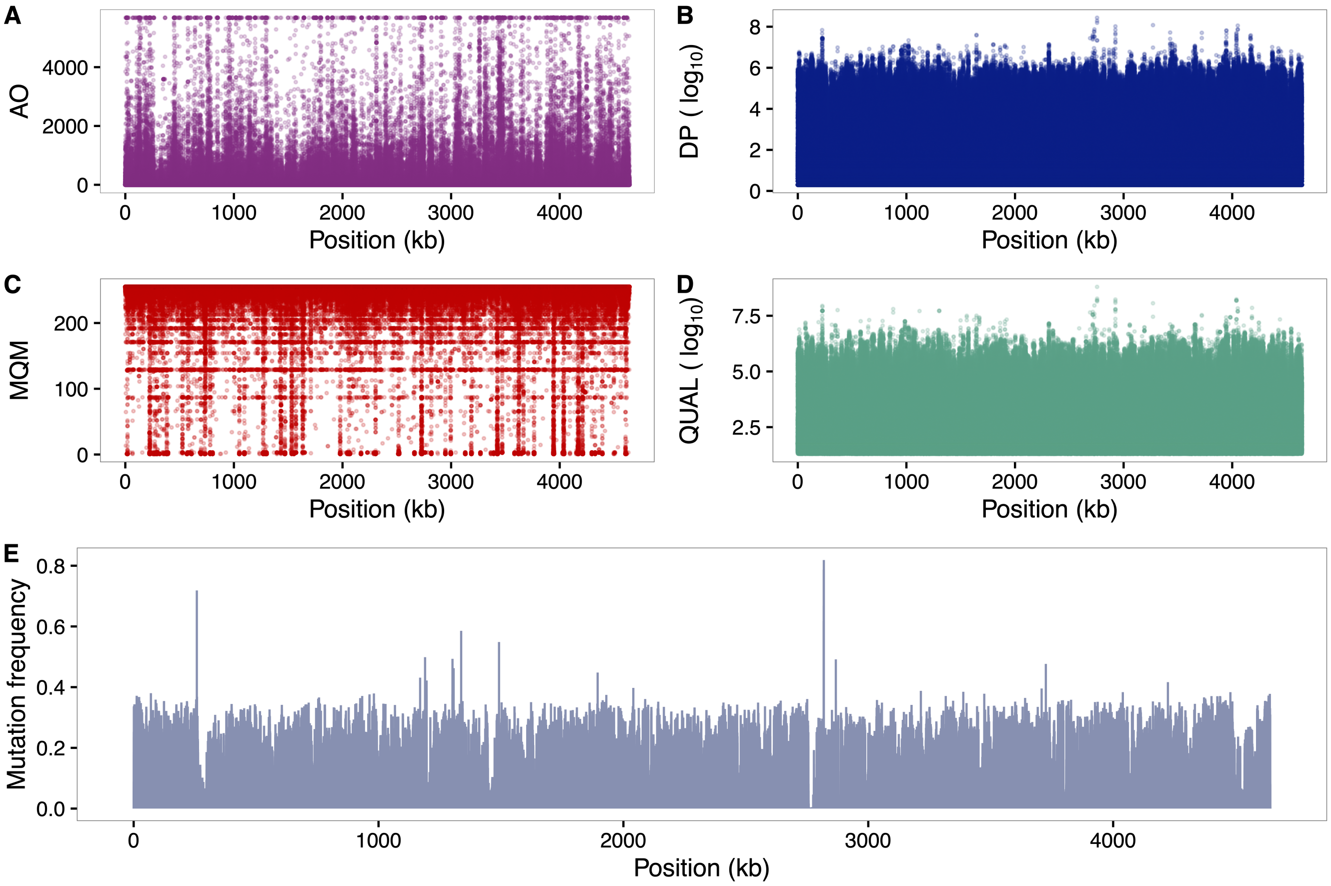


**Figure 6: Summary QC of synonymous variants identified across the *E. coli* genome.** VCF data filtered upon **(A)** alternate allele observation count (AO), with counts thresholded at the 99.9th percentile of the distribution of counts, **(B)** read depth (DP, $\log_{10}$-scaled), **(C)** mean mapping quality of observed alternate alleles (MQM), **(D)** Phred-scaled quality (QUAL, $\log_{10}$-scaled), and **(E)** frequency of mutations per sample across loci, determined by filtering the total number of alternate alleles in called genotypes (AC) and normalising by total number of samples.

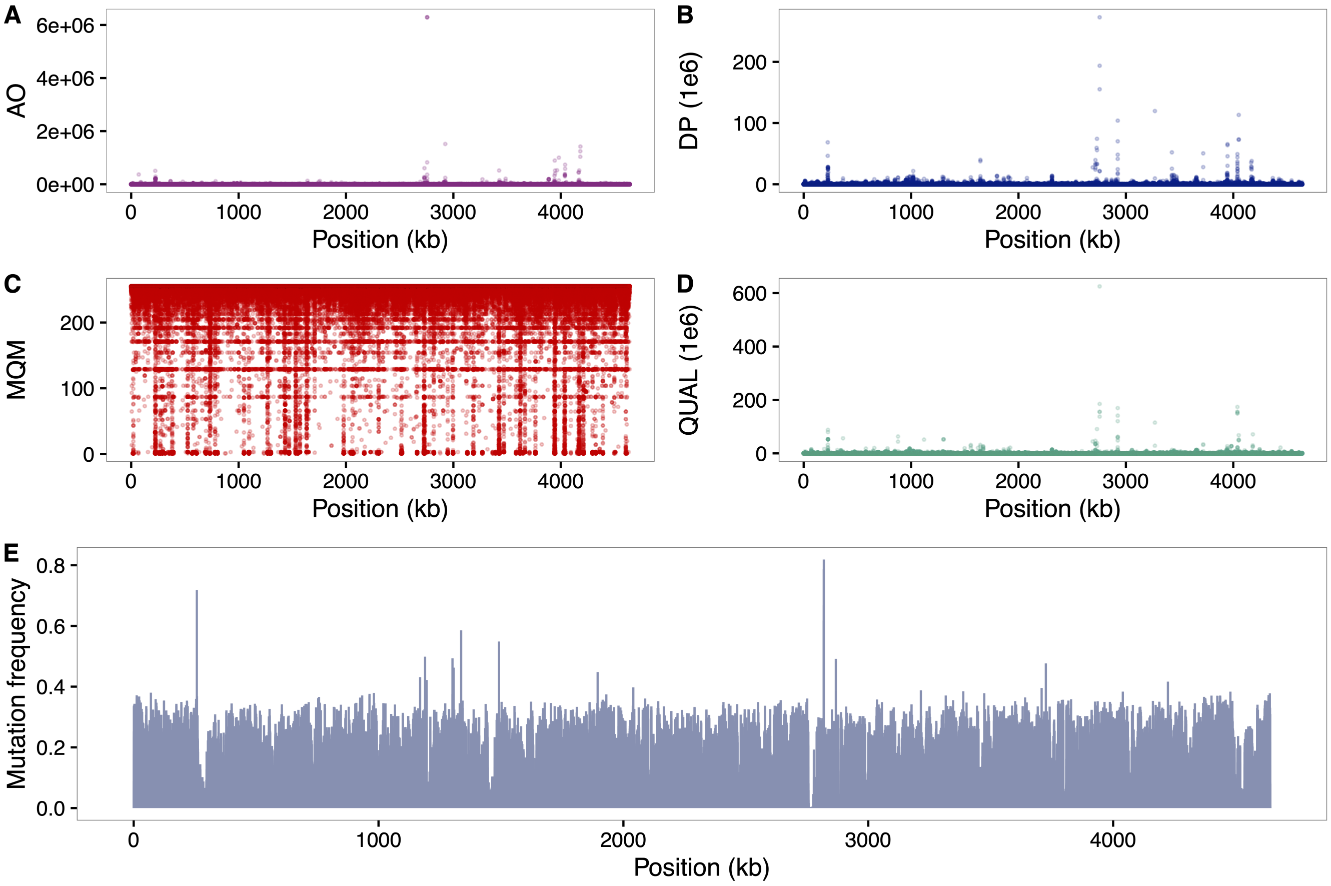


**Figure 7: Variant QC using raw values.**

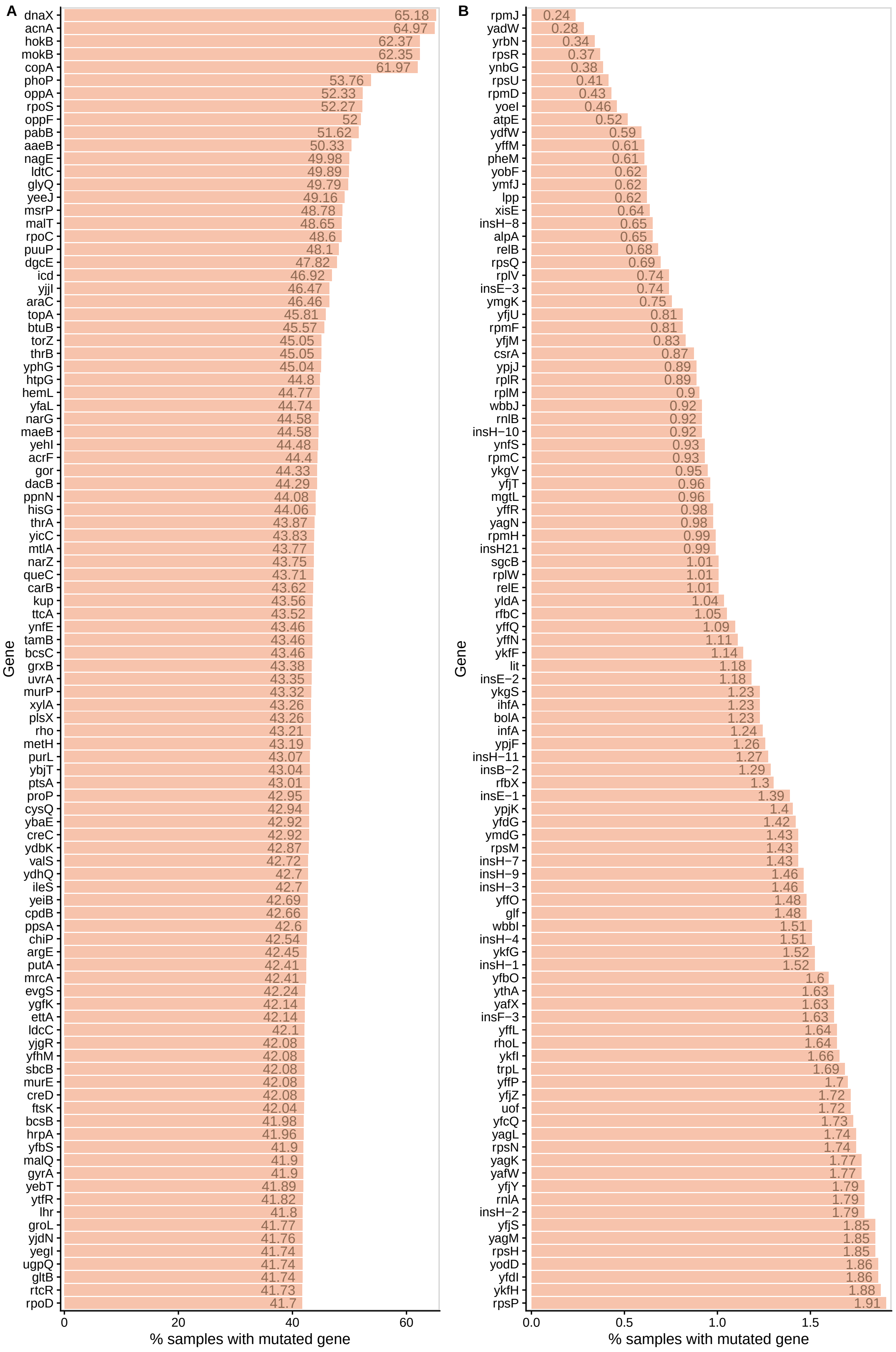


**Figure 8: Top 100 mutated genes.**

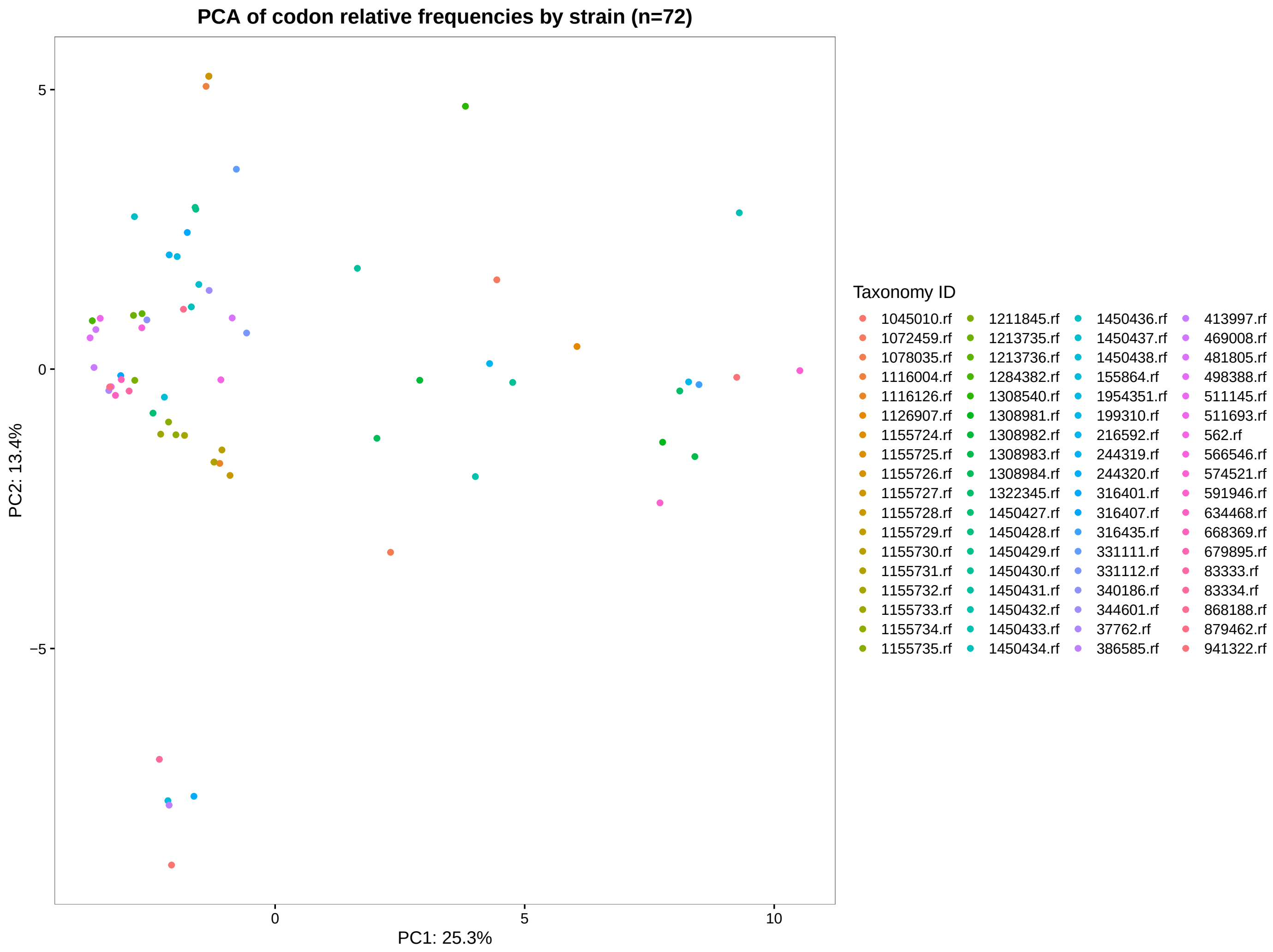


**Figure 9: PCA of codon usage per strain.**

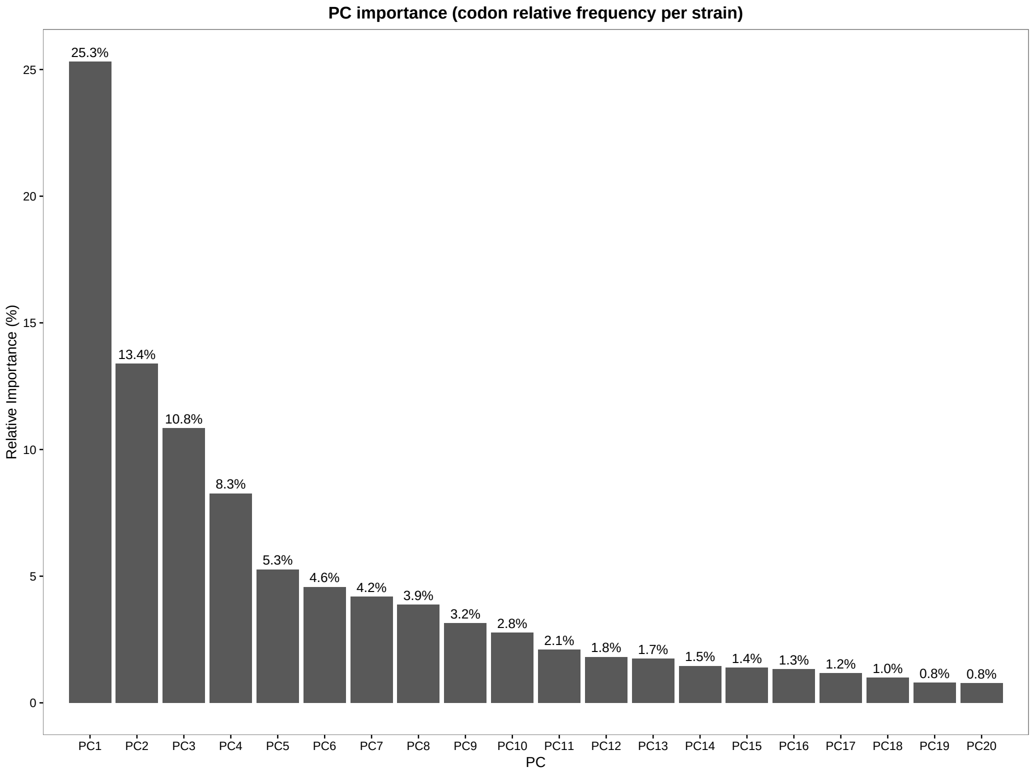


**Figure 10: Strain PC scores.**

**Figure 11: Box plots comparing log2 distribution of relative codon frequency difference in *CUBseq* and Kazusa *E. coli* K-12 genes.** Each box represents the absolute difference in relative codon frequencies, grouped into amino acids, and coloured by its corresponding amino acid property (STOP codons are coloured grey). Comparison of codon frequencies in *CUBseq* highly expressed genes (CUBseq-HEG) **(A)** and *CUBseq* lowly expressed genes (CUBseq-LEG) **(B)**, with Kazusa genes. **(C)** Comparison of codon frequencies in CUBseq-LEG with CUBseq-HEG. Boxplots show the median (thick centre line), along with interquartile ranges (box, $25\%$ to $75\%$) and whiskers extending $1.5$x the interquartile range.

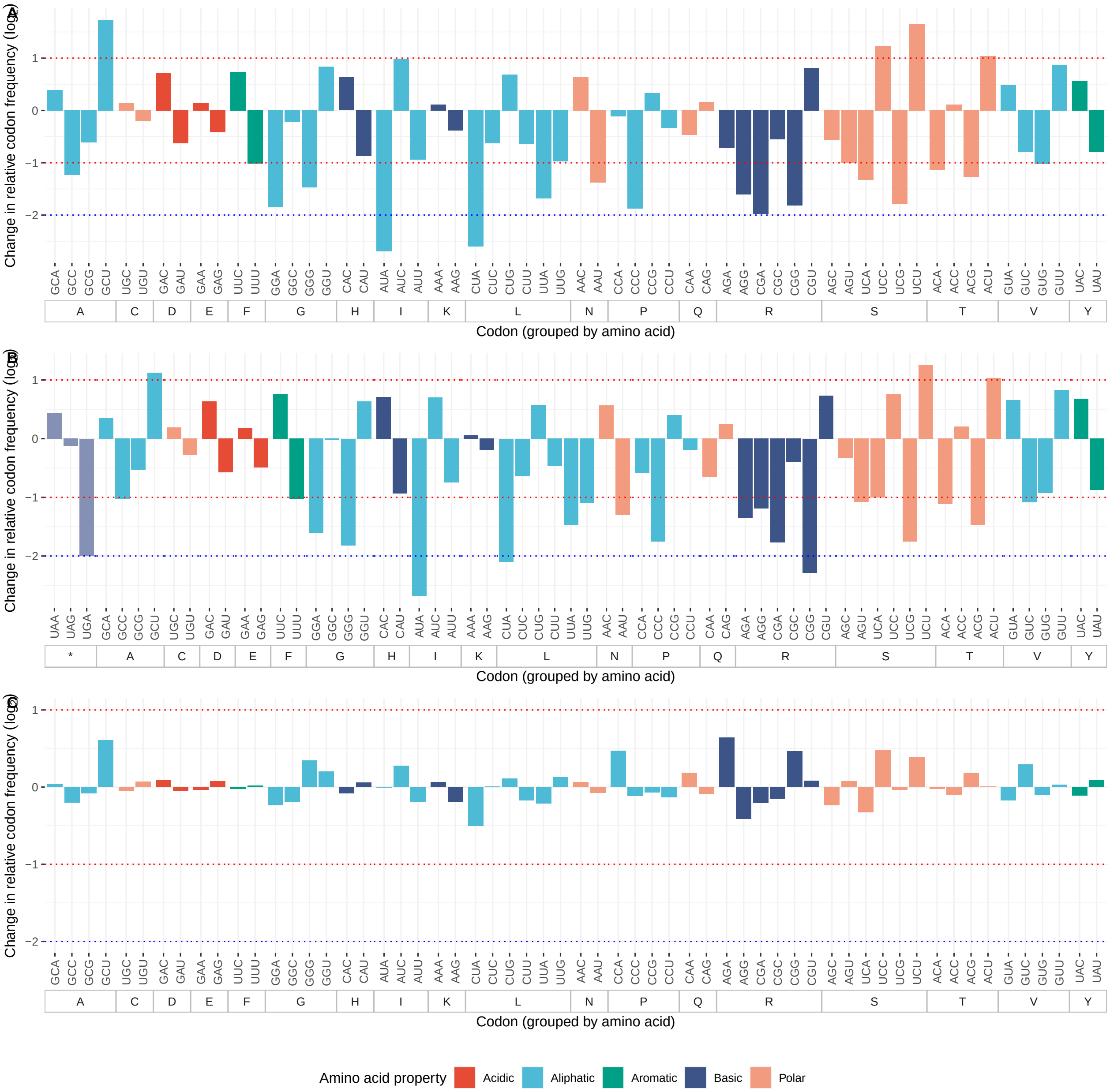


**Figure 12: Bar plots comparing relative codon frequencies in cubseq and Kazusa *E. coli* K-12 genes.** Each bar represents the absolute difference in codon frequency grouped into amino acids, and coloured by its corresponding amino acid property (STOP codons are coloured grey). Comparison of codon frequencies in cubseq highly expressed genes (cubseq-HEG) **(A)** and cubseq lowly expressed genes (cubseq-LEG) **(B)**, with Kazusa genes. **(C)** Comparison of codon frequencies in cubseq-LEG with cubseq-HEG.

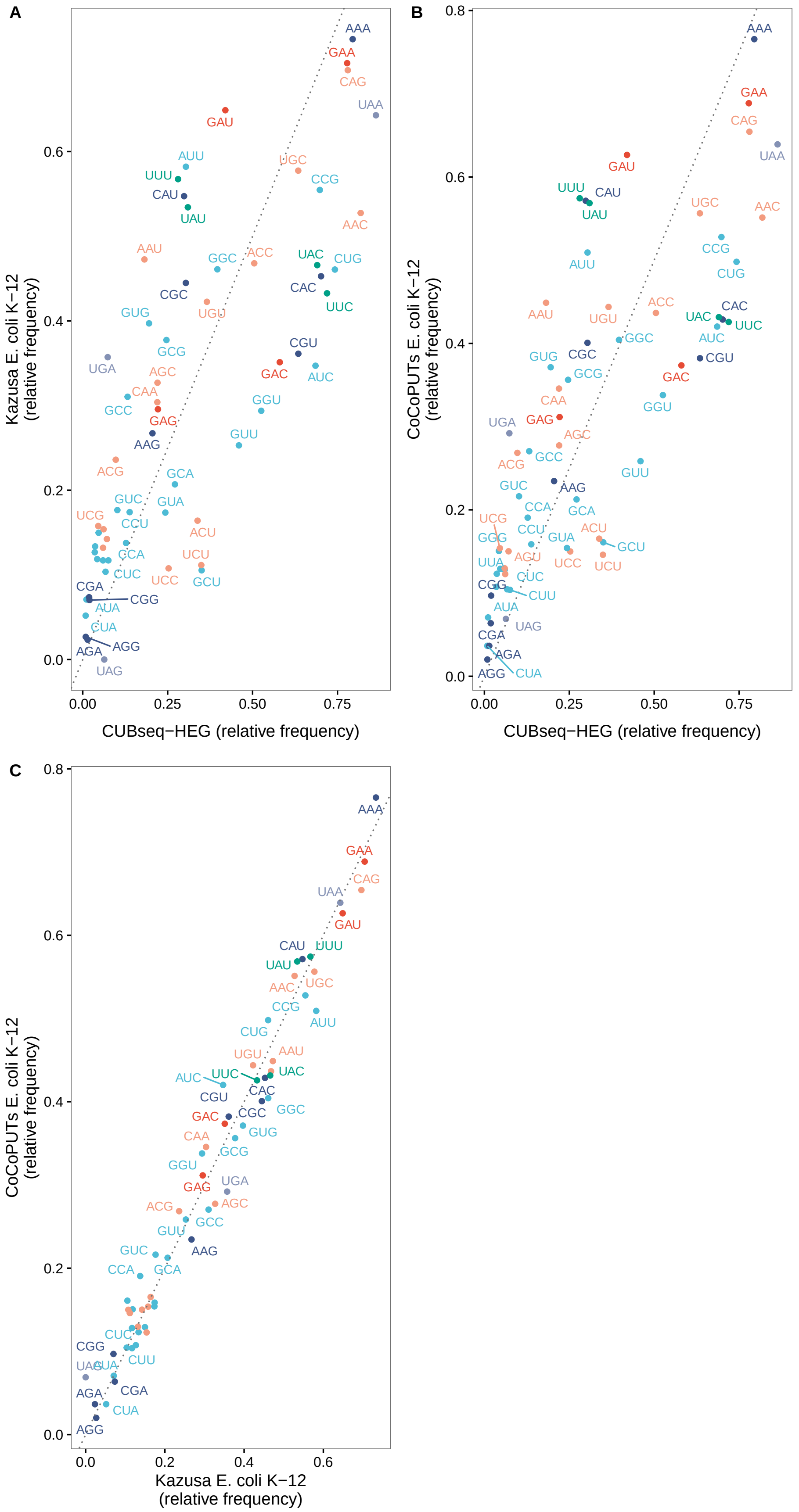


**Figure 13: Comparison of relative codon frequencies in CUBseq and Kazusa *E. coli* K-12 genes.** Scatter plots comparing codon frequencies in Kazusa *E. coli* K-12 versus cubseq **(A)** highly expressed (cubseq-HEG) and **(B)** lowly expressed genes (cubseq-LEG). **(C)** Comparison of codon frequencies in cubseq-LEG with cubseq-HEG. Each point represents a codon and its relative frequency, labelled by its corresponding amino acid and coloured by its corresponding amino acid property (STOP codons are coloured grey). If there were no differences present between cubseq and

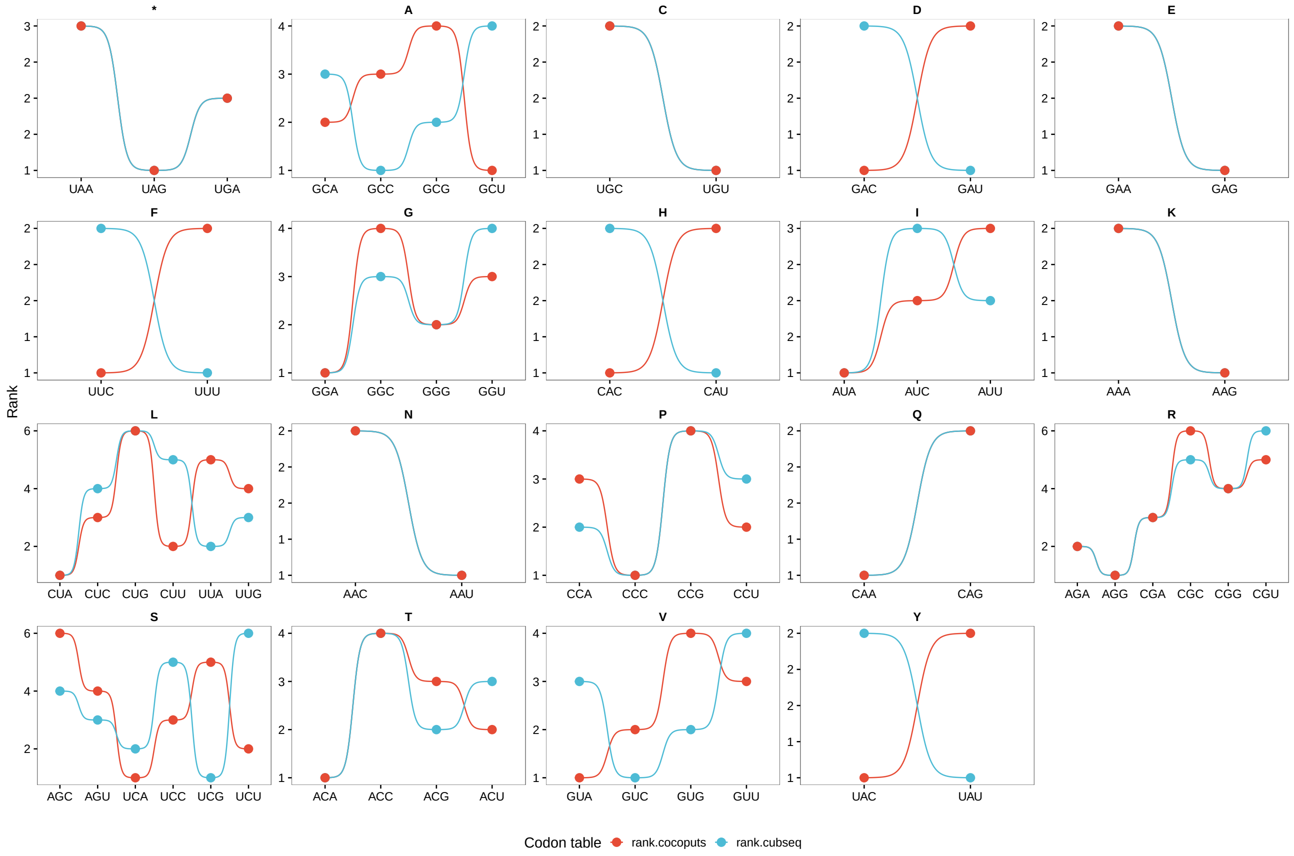


**Figure 14: Codon rank analysis.**
